# Cold shock as a screen for genes involved in cold acclimatization in *Neurospora crassa*

**DOI:** 10.1101/259580

**Authors:** Michael K. Watters, Victor Manzanilla, Holly Howell, Alexander Mehreteab, Erik Rose, Nicole Walters, Nicholas Seitz, Jacob Nava, Sienna Kekelik, Laura Knuth, Brianna Scivinsky

## Abstract

When subjected to rapid drops of temperature (cold shock), Neurospora responds with a dramatic, but temporary shift in its branching pattern. While the cold shock response has been described morphologically, it has yet to be examined genetically. This project aims to begin the genetic characterization of the cold shock response and the associated acclimatization to cold environments. We report here the results of a screen of mutants from the Neurospora knockout library for alterations in their morphological response to cold shock and thus, their ability to acclimatize to the cold. Three groups of knockouts were selected to be subject to this screen: genes previously suspected to be involved in hyphal development as well as knockouts resulting in morphological changes; transcription factors; and genes homologous to E. coli genes known to alter their expression in response to cold shock. Several strains were identified with altered responses. The genes impacted in these mutants are listed and discussed. A significant percentage (81%) of the knockouts of genes homologous to those previously identified in E. coli showed altered cold shock responses in Neurospora – suggesting that the response in these two organisms is largely shared in common.

---

The environmental conditions that life must contend with can vary widely. Organisms have evolved a wide range of mechanisms for contending with these changing conditions. For the filamentous fungus Neurospora, growth continues through nearly the entire range of temperatures (above freezing) that is observed in this environment. Although the rate of tip extension varies linearly with temperature (Watters et al 2000), the branch density (the statistical distribution of distances between branch sites along a linear growing hypha) remains constant across this range (Watters *et al.* 2000) allowing the fungus to continue to infiltrate its environment at the same density. Temperatures progressing through this range would be expected to have dramatic impacts on enzyme activity generally (and thus overall metabolism), but also directly on features critical to growth such as membrane fluidity, DNA/RNA stability and the rates of transcription and translation.

## Response to cold shock

When the apparent independence of branching and temperature is tested by rapid temperature shifts, a 3-phase response is observed (Figure 1, Watters *et al.* 2000, Watters 2013). The initial response to cold shock is the growth of a single longer than normal unbranched segment. This was termed the “Lag” phase of the response. This phase is followed by a series of closely spaced apical branch points, termed the “Apical” phase. Apical branch formation has been previously associated with the disruption and attempted reorganization of the normal tip-growth apparatus (Reynaga-Peña *et al.* 1995, Riquelme & Bartnicki-Garcia 2004), a mechanism distinct from that thought to be involved in lateral branching. Finally, with continued incubation at the lower temperature, the colony returns to lateral branching, termed the “Recovery” phase. Growth in this phase of the response resembles that which would be seen had the colony been grown at 4°C continuously. The same density is observed for growth subjected to constant incubation at 4°C (or any other fixed temperature) as well (Watters et al 2000). Thus, the cold shock response appears to be a temporary disturbance to a homeostatic system which maintains branch density at a constant, evolutionarily favored, value.

**Figure 1:**
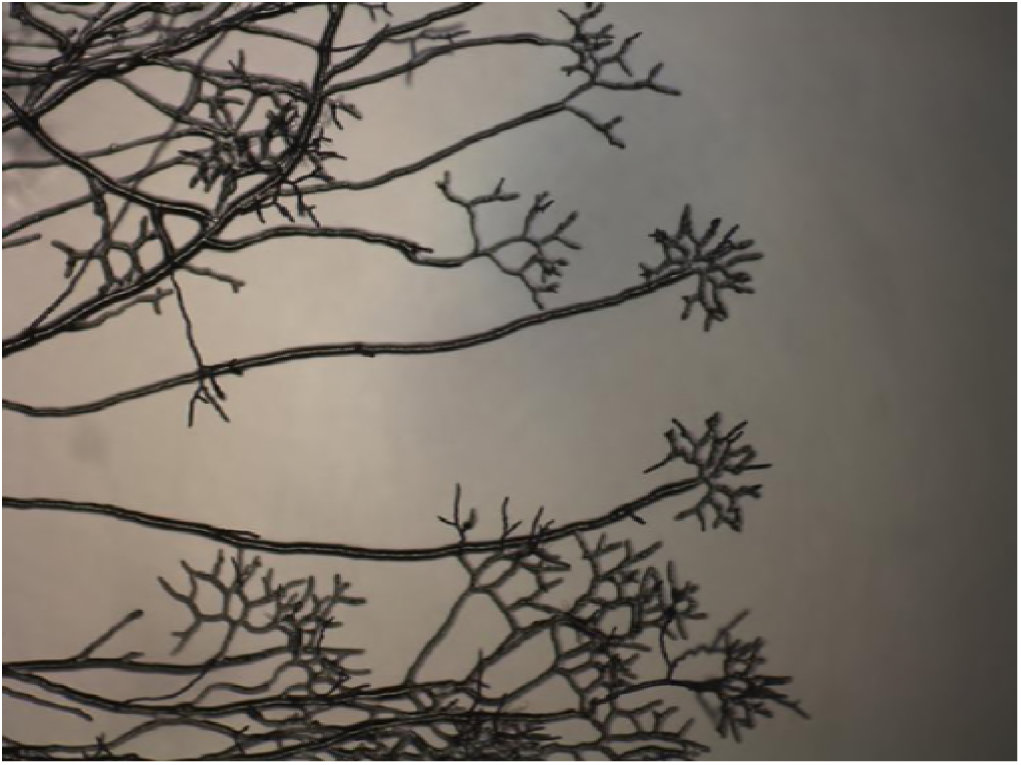
Cold shock response in wild-type Neurospora:

Thus, Neurospora appears to incorporate a system which has the effect of maintaining homeostasis for critical cellular characteristics. At least one impact of this system is that branch density is maintained at widely different temperatures. The morphological effects of cold shock are the indirect consequence of this system’s staged process of adjusting cellular conditions in order to compensate for the new growth temperature.

## Cold shock response of bacteria

Homeostasis in the face of temperature changes and more specifically the response to cold shock has been extensively studied in bacterial systems for over 20 years. The effect of cold shock is manifest in multiple cellular systems including: membrane rigidity (Shivaji & Prakash 2010), stability of secondary structures in DNA/RNA (Phadtare 2004), efficiency of protein folding (Phadtare 2004) and ribosome function (Gualerzi *et al.* 2011). While much remains to be described in these systems, cold shock appears to result in a multi-stage response (Phadtare 2004). First, a lag period in which growth and translation of proteins generally cease. This is followed by an adjustment phase in which specific cold-shock proteins which compensate for the changes brought on by the cold are preferentially translated (Giuliodori *et al.* 2004). In the final stage, growth continues otherwise normally, but at a reduced rate. DNA microarray transcription profiling of the cold shock response in E. coli by Phadtare and Inouye (2004) has shown that several hundred genes respond to cold shock, either being transiently induced/repressed or showing prolonged induction/repression. Analogous responses to cold shock and/or cold acclimation have been observed in diverse organisms including plants (Guy 1999) and animals (Canclini & Esteves 2007). Attempts to uncover cold shock proteins in fungi (Fang & Leger 2010) have met with mixed success.

## Connections between cold shock in bacteria and Neurospora

It is tempting to draw parallels between what is known about cold shock in bacterial systems and the observed response of Neurospora to similar cold shocks. Many of the systems affected during bacterial cold shock would be expected to impact fungal tip growth and branching (most obviously, membrane fluidity). Beyond that however, the nature and timing of the two responses are similar. Both responses can be adjusted by changing the intensity of the cold shock with more mild shocks (lower temperature differences) producing more mild responses and more severe shocks (larger temperature differences) producing more severe responses. Furthermore, the dynamics of the responses parallel each other. In each, there is a multistage response. There is an initial response which is transient in nature, followed by a more long-term response which largely represents a return to normal growth.

## Guiding Hypotheses

The hypothesis of this project was that the observed cold shock response of Neurospora is a consequence of a cellular response homologous to that induced by cold shock in bacteria. Under this hypothesis, the observed, transient morphological changes are an aspect of the primary response which is that of the fungal cell adjusting itself to growth in the cold via a manner which is shared in common with simpler organisms. This hypothesis was tested by screening Neurospora knockout strains impacting genes homologous to those identified in E. coli which alter their expression patterns in response to cold shock. Of 68 knockouts screened, 55 (81%) showed changes in the morphological response to cold shock, suggesting a strong connection between the responses of E. coli and Neurospora to cold shock.

Additionally, Neurospora strains with knockouts of known transcription factors, genes impacting morphology, or genes homologous to those in yeast known to affect polar growth were similarly screened. In this screen, of 357 knockouts examined, 69 (19%) showed changes in the response to cold shock. Together, these screens provide the first molecular underpinning to the cold shock response in Neurospora,

## MATERIALS AND METHODS

### The Neurospora targeted deletion collection

As part of the Neurospora Genome Project, a collection of strains containing disruptions in presumptive genes has been constructed (Colot *et al.* 2006). Strains representing deletions of most of the genes of the Neurospora genome are available from the Fungal Genetics Stock Center (McCluskey 2003). As each deletion strain has been altered in a single, previously identified, presumptive gene – going from phenotype to sequence is greatly simplified.

The accession numbers listed in Table 1 represent the locus number of the gene subject to inactivation in the knockout strain under test. Every annotated gene in *Neurospora crassa* has been assigned a locus number of the form NCU####. The functions reported (Tables 1 – 3) are those associated with the genes as annotated on the FungiDB database as of July 2017: fungidb.org/fungidb/. The reported functions are based solely on the annotations currently associated with those strains and have not been independently confirmed by the authors of this study. The genes identified in this study have been annotated on FungiDB to reflect these results.

**Table 1:**
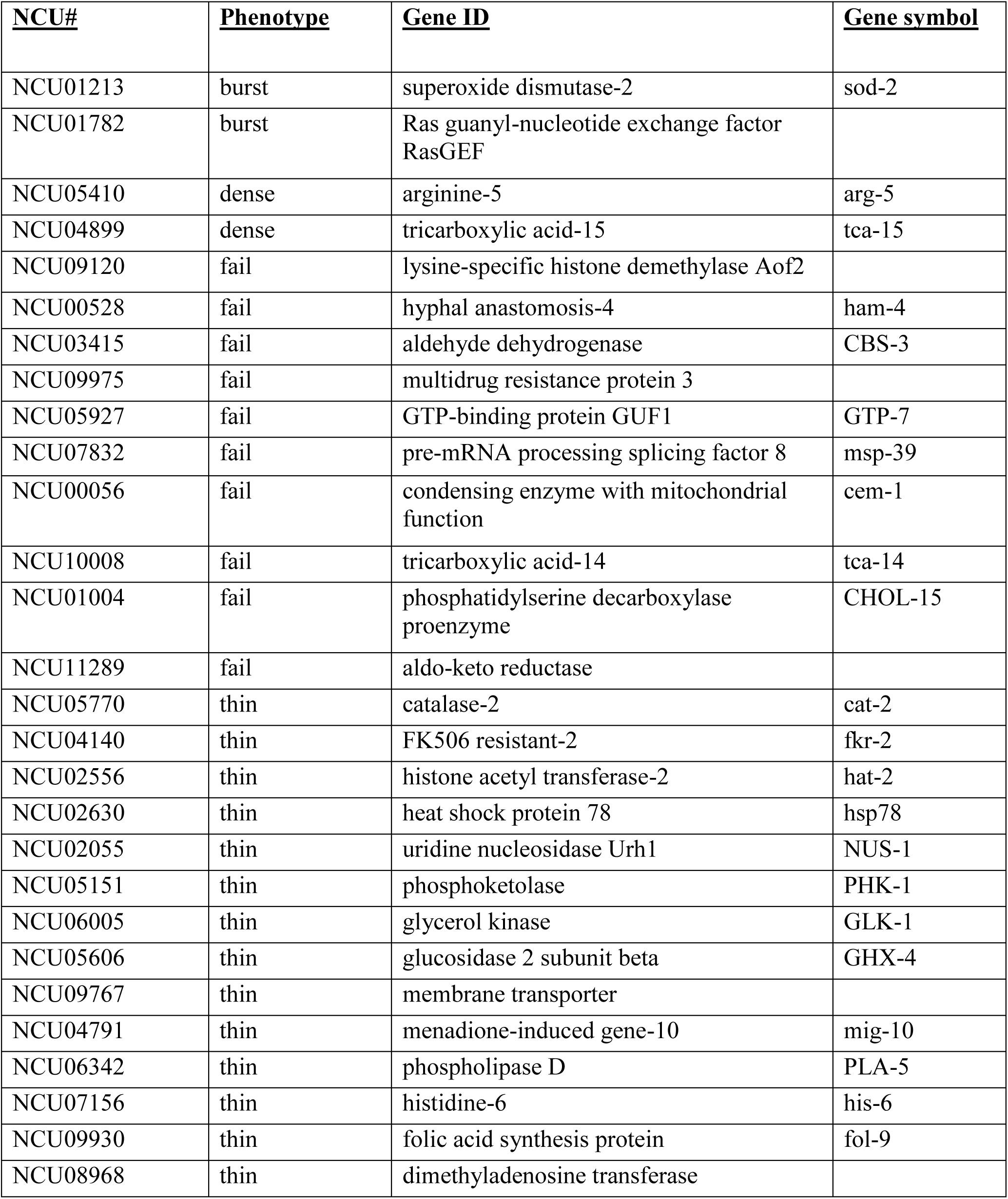
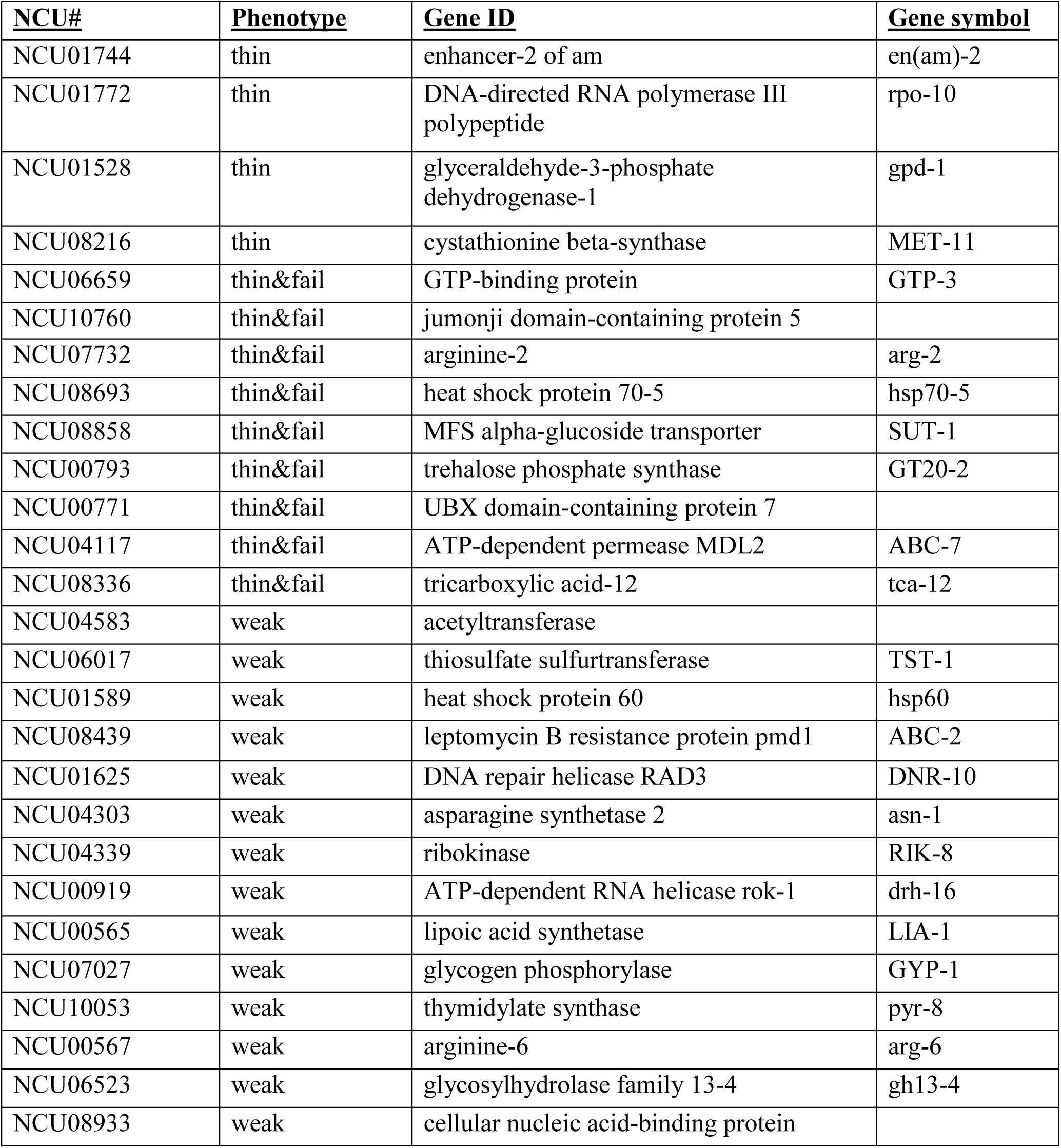
Results of screening 68 knockouts selected as orthologs of genes known to alter expression in response to cold shock in E coli:

**Table 2:**
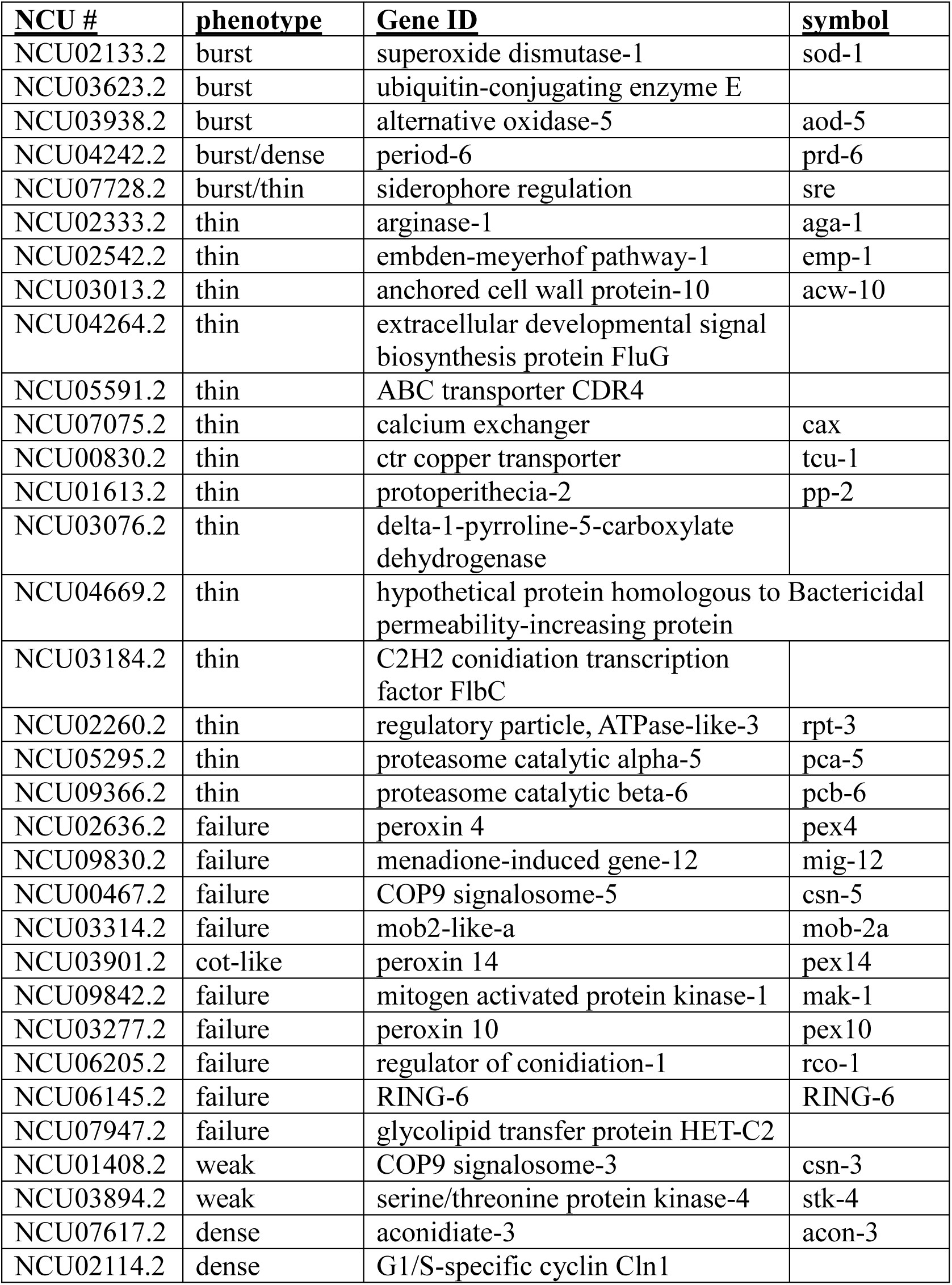
Results of screening 148 knockouts on the “morphological” and “hyphal growth” plates:

**Table 3:** Results of screening 208 knockouts of Neurospora transcription factors:

| NCU# | Phenotype | Gene ID | Gene symbol |
| --- | --- | --- | --- |
| NCU05051 | thin | COL-23 | col-23 |
| NCU01629 | thin | hypothetical protein |  |
| NCU04561 | thin | melanization defective-1 | mld-1 |
| NCU05909 | weak | hypothetical protein |  |
| NCU00499 | weak | all development altered-1 | ada-1 |
| NCU07945 | weak | fungal specific transcription factor | tah-4 |
| NCU08658 | weak | zinc finger transcription factor-50 | znf-50 |
| NCU02017 | fail | CBF/NF-Y family transcription factor | ada-2 |
| NCU00097 | fail | BEAK-1 | bek-1 |
| NCU07561 | fail | hypothetical protein |  |
| NCU00499 | fail | all development altered-1 | ada-1 |
| NCU02173 | fail | zinc finger transcription factor-52 | znf-52 |
| NCU02356 | fail | white collar 1 | wc-1 |
| NCU08294 | fail | nitrogen assimilation transcription factor nit-4 | nit-4 |
| NCU08294 | fail | nitrogen assimilation transcription factor nit-4 | nit-4 |
| NCU06990 | fail | hypothetical protein |  |
| NCU07788 | fail | fungal specific transcription factor | col-26 |
| NCU06028 | fail | quinic acid utilization activator | qa-1F |
| NCU07945 | fail | fungal specific transcription factor | tah-4 |
| NCU06068 | fail | fungal specific transcription factor | col-25 |
| NCU02214 | fail | TAH-2 | tah-2 |
| NCU03070 | burst | hypothetical protein |  |
| NCU03962 | dense | hypothetical protein |  |
| NCU03356 | dense | hypothetical protein |  |
| NCU03905 | dense | hypothetical protein |  |
| NCU01154 | dense | submerged protoperithecia-1 | sub-1 |
| NCU00144 | dense | hypothetical protein |  |
| NCU03417 | dense | hypothetical protein |  |
| NCU06990 | dense | hypothetical protein |  |
| NCU03120 | dense | hypothetical protein |  |
| NCU08000 | thin & fail | cutinase transcription factor 1 alpha | far1 |
| NCU08651 | thin & fail | zinc binuclear cluster-type protein | col-27 |
| NCU05536 | thin & fail | hypothetical protein |  |
| NCU07705 | thin & fail | C6 finger domain-containing protein | clr-1 |

### Selection of knockout strains to screen

A screen of the entire library was determined to be impractical. We instead screened an abbreviated subsection of the library chosen to be more likely to yield positive responses.

First, knockouts of genes homologous to those which show altered transcription in E. coli subjected to cold shock. The protein sequences of E. coli genes identified by (Phadtare and Inouye 2004) were retrieved from the E. coli database (ecocyc.org/). These amino acid sequences were then fed into a BLAST search on the NIH NCBI site (blast.ncbi.nlm.nih.gov/Blast.cgi) with the output limited to Neurospora sequences in order to identify their nearest Neurospora homologs. These homologs were then searched on FungiDB to determine which had knockout strains available. From this final list, 68 were selected randomly for screening in this study. This set was selected to determine the degree of relationship between the cold shock response in E. coli and Neurospora.

Second, two sets of knockouts from the library (hyphal growth and morphological) were included in this screen. One plate (identified as “plate 29 – morphologicals” by the FGSC) contained 71 strains with knockouts known to cause morphological changes. The second plate (identified as “Hyphal Growth Set” by the FGSC) contained 78 strains with knockouts in genes homologous to genes in yeast known to affect polar growth.

Lastly, knockouts of known transcription factors in Neurospora were selected for cold shock screening. This collection is available as a set from the Fungal Genetics Stock Center (McCluskey 2003). It was selected for this screen to determine which transcription factors play a role in signaling to the cell that cold adaptation genes must be activated.

### Media

Media and culturing procedures were those described in Davis & deSerres (1970). Growth described as being on “minimal” was on Vogel’s minimal medium (Davis & deSerres 1970).

### Screen

The selected knockout strains were subjected to a screen looking for altered responses to cold shock. Wild-type Neurospora progresses through a three-stage response following a shift into the cold. As detailed above, these are the Lag, Apical and Recovery phases. As the cold shock response is known to be stronger following more dramatic temperature shifts (Watters et al. 2000) we initially grew strains at 33°C and shifted to 4°C. Strains were inoculated onto Vogel’s Minimal Medium and incubated overnight at 33°C. The next morning plates were moved to 4°C. After an overnight incubation at 4°C, the strain’s response to cold shock was photographed and evaluated. Variations in the cold shock response from that of wild-type Neurospora were judged qualitatively and based primarily on the morphology of the “apical” phase of branching.

### Photomicroscopy

Growing cultures were examined and photographed using a Motic 10MP digital camera attached to a Wolfe Beta Elite trinocular microscope. Photographs were taken of well separated, leading hyphae.

## RESULTS AND DISCUSSION

During the initial study of the cold shock response in Neurospora (Watters et al 2000) it was observed that several classical morphological mutants (most notably “granular” and “delicate” produced altered responses to cold shock, demonstrating that mutants could be obtained which influenced this process. We chose to screen mutants from the Neurospora knockout library for their cold shock response in order to provide a genetic grounding to this process which has, thus far, been lacking. We chose to use the mutants of the knockout library instead of the products of a random mutagenesis as the knockouts allow an immediate identification of gene function in most cases.

Knockout strains displaying an altered morphological response to cold shock were classified according to the specific variation they displayed. Examples are shown in Figure 2. The “burst” phenotype was defined as displaying a large number of growing tips which stop growing, swell and then structurally fail leaving a pool of cytoplasm at the tip. The “fail” phenotype was defined as failing to display the apical branch phase characteristic of cold shock. In the “fail” response, growth proceeds normally with lateral branching following cold shock. The “thin” phenotype was defined by a very rapid decrease in hyphal diameter following cold shock. It was common to observe “thin” in combination with other altered cold shock responses. The “dense” phenotype was defined by displaying apical branching with visibly shorter distances between branch points following cold shock relative to the response in wild-type. The “weak” phenotype was defined as the opposite – an apical branch phase with visibly longer distances between branch points relative to wild-type following cold shock. Finally, the “cot-like” phenotype was characterized by a lack of apical branching, but a shift to tightly spaced lateral branches which morphologically resembled the growth of the traditional *cot* mutants at the restrictive temperature.

**Figure 2:**
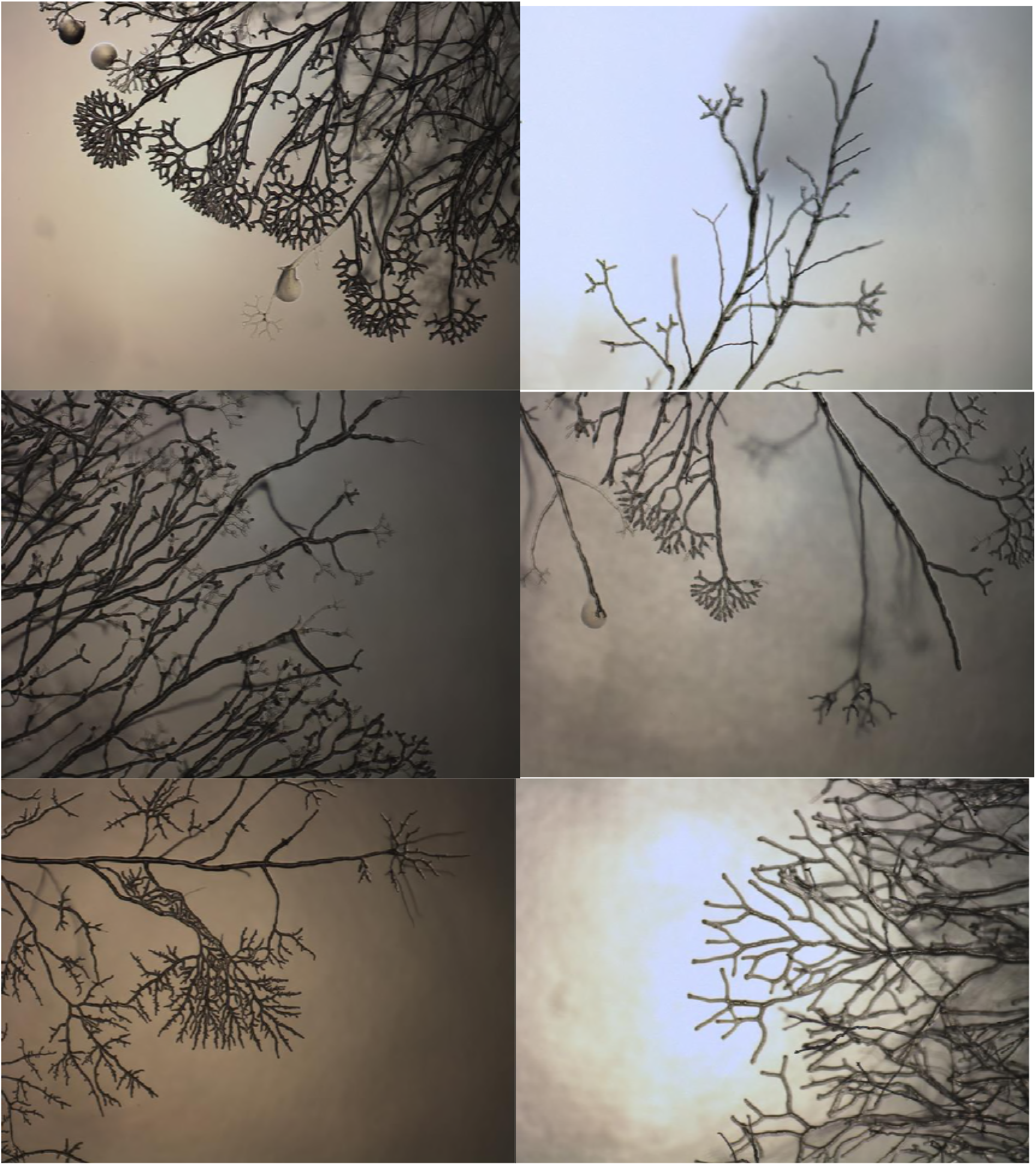
Alternate cold shock morphologies displayed: Bursting tips, failure to cold shock, shift to thin hypha, extra dense apical branching, cot-like, weaker than normal apical branching

We can imagine two distinct groups of genes to be identified by these screens. The first would be genes directly involved in cold adaptation which are responsible for altering the cell to accommodate the altered environment. This first group of functions, we could very well expect to be generalized to a wide variety of organisms. The second would be genes coding for proteins which are individually temperature sensitive and compensate for temperature changes by either altering regulation to compensate for the change in activity or shifting activity to a paralog which functions better at the new temperature. This second group we might expect to be more species specific.

### Screen of E. coli cold shock gene homolog knockouts

A total of 68 Neurospora strains with knockouts of genes homologous to E. coli genes which alter transcription in response to cold shock (Phadtare and Inouye 2004) were screened. A total of 55 (81%) showed altered morphology to cold shock (Table 1). The knockouts displaying altered response to cold shock represent a variety of cellular functions. There does not to be any correspondence between the response of a gene in Ecoli (up regulated, down regulated, transient or sustained), or its function (membrane metabolism, ROS control etc) (Phadtare and Inouye 2004) and the morphology displayed during cold shock.

The screen of cold shock orthologs provides a test of the hypothesis that the cold shock response in both E. coli and Neurospora share a great deal of their cold shock response in common. The very high percentage of overlap between genes playing a role in these two widely separated organisms argues that the two responses are functionally related to a large degree. The large fraction of overlap also argues that the majority of the genes identified in these two organisms are involved generally in cold adaptation. The results also suggest that screening for deviations from the normal cold shock response morphology is an effective tool for detecting genes important in cold adaptation.

No correlation was observed between the transcription change observed in E. coli (Phadtare and Inouye 2004) and the observed cold shock phenotype observed in the knockout strain of its Neurospora ortholog. Similarly, no correlation was observed between the cold shock phenotype observed and the annotated function of the genes affected in these strains.

### Screen of Morphological/Hyphal plates

A total of 149 selected mutant strains from the Neurospora knockout library were previously segregated into two collections. The “Morphological” collection resulted in known morphological variations in the knockout strains. The “Hyphal” collection consisted of knockouts of genes previously suspected to play a role in hyphal growth. These two collections were screened for alterations to their response to cold shock. In total, 35 (23%) strains were identified (Table 2) that displayed variant cold shock responses. The altered responses fell into several phenotypic categories. The genes impacted in each knockout strain are identified in Table 2.

As with the E. coli orthologs, the genes identified from this collection which displayed altered response to cold shock appear to be involved in a number of different cellular functions. In this case they include lipid/membrane metabolism, protein degradation/turnover, gene regulation, protein regulation, and reactive oxygen control.

The morphological/hyphal knockouts were previously screened for temperature-dependent branch density (Watters et al. 2011). Comparing the strains identified above with alterations to their cold shock response to those previously determined to show temperature-dependent branching we find only a modest overlap with the following strains showing altered phenotypes in both: NCU02333, NCU00830, NCU04242, NCU02114, NCU04264, and NCU03076. Examining the overlap statistically via Chi-square (calculations not shown) yields a p value greater than 0.9, strongly suggesting that the overlap is random. This suggests that these two screens (cold shock vs temperature sensitive branching during steady-state growth) are independent. This leads us to conclude that the cold shock response and temperature-dependent branching are independent aspects of cold adaptation, highlighting the different genes involved in short-term adaptation to the cold as opposed to those required for sustained growth in cold environments. Additional screens of the knockout library for strains displaying growth rate dependent branching, and comparing them to those with an altered cold shock response will allow us to further examine the apparent independence of these two morphological screens.

### Screen of transcription factor knockouts

A total of 208 Neurospora strains with knockouts in genes which function as transcription factors were screened for their response to cold shock. In all, 34 (16%) showed altered morphology to cold shock (Table 3).

The transcription factor screen identifies a number of genes which may function in a broad way to regulate multiple genes to the purpose of cold adaptation.

As with the knockouts of orthologs of E. coli cold shock responding genes, the mutant strains identified in the additional screens show no observed correlations between the phenotypes observed and the annotated functions of the genes with a variety of functions being associated with the observed cold shock variations.

In conclusion, the gene functions highlighted by these screens (Table 1-3) are diverse. It is unclear how the diverse gene network, partially exposed here, coordinates for the function of temperature acclimatization. The results presented here demonstrate a strong relationship between the cold shock responses of E. coli and *Neurospora crassa*. The phenotype under examination here (morphological response to cold shock) appears to be influenced by a diverse network of genes. Similar diversity of function has been observed in other examinations of morphogenesis in Neurospora (Seiler & Plamann 2003). Further work on cold acclimatization, including a broad survey of the full Neurospora knockout library should help clarify these connections.

## ACKNOWLEDGEMENTS

This work was supported by a grant from the Indiana Space Grant Consortium (INSGC). We would also like to thank Kevin McCluskey and everyone at the Fungal Genetics Stock Center for supplying the knockout strains and their diligent work in support of the fungal genetics community over the years.

## Introduction

In both Neurospora and *E. coli*, there is a multistage response to cold shock. There is an initial response which is transient in nature, followed by a more long-term response which largely represents a return to normal growth. Neurospora grows via extension at a hyphal tip with periodic branching which is typically lateral (Figure 1A). However, when Neurospora is subjected to cold shock, a multi-phase morphological response is observed (Figure 1B, Watters *et al.* 2000, Watters 2013). The morphological effects of cold shock are the indirect consequence of this system’s staged process of adjusting cellular conditions in order to compensate for the new growth temperature.

**Figure 1:**
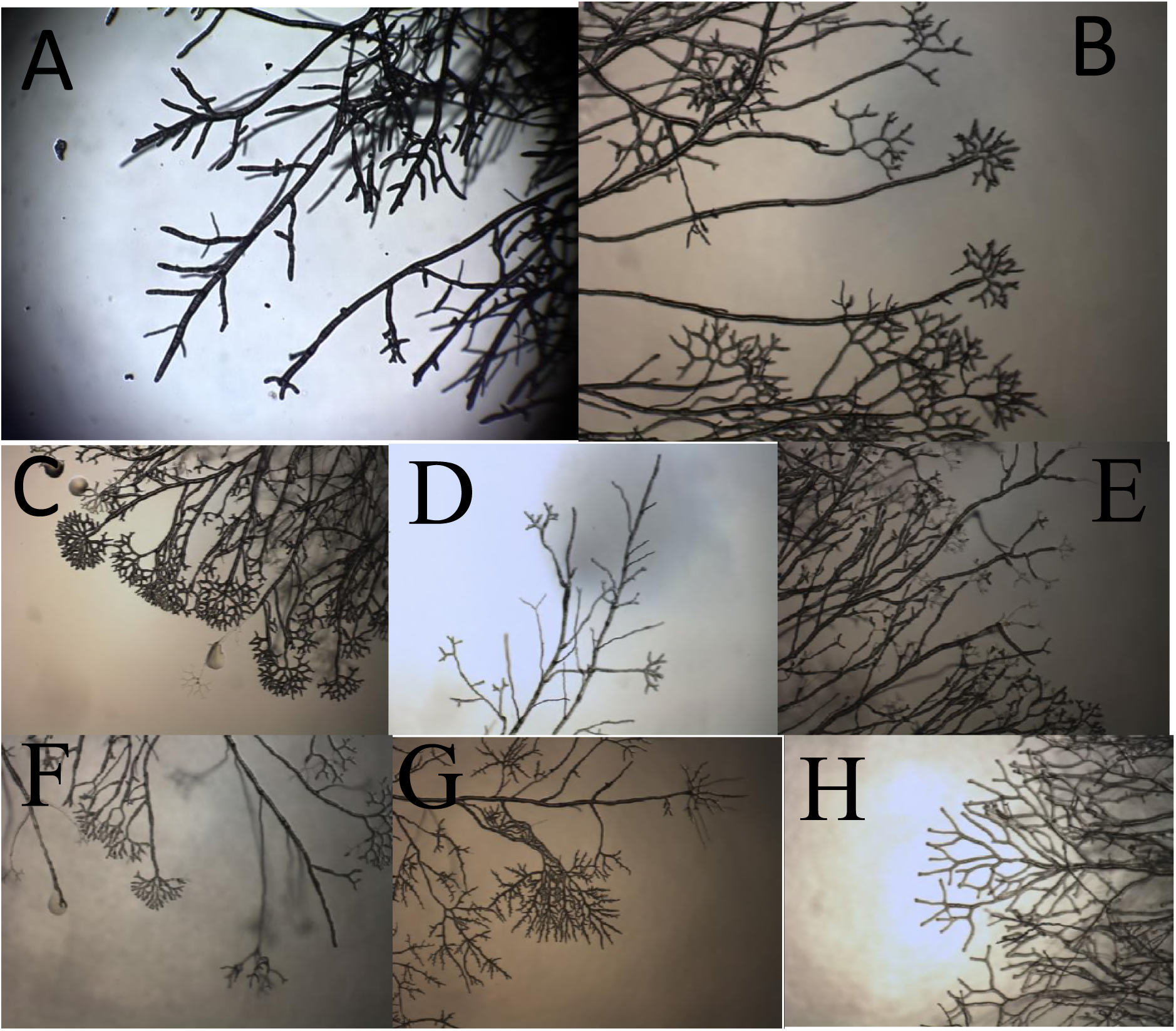
Conventional growth vs cold shock in wild-type and mutant Neurospora: A) Wild-type (Oak Ridge) Neurospora growth at 33⁰C, B) cold shock response in wild-type Neurospora, While many of the knockout strains tested displayed a morphological response to cold shock indistinguishable from that of wild-type, alternative morphologies were observed. These were classified into categories, examples of which are shown here. Examples of the alternate cold shock phenotypes displayed with the identity of the mutant shown as the example are shown: C) Burst: tips of growing hypha burst commonly (NCU02133, superoxide dismutase-1), D) Fail: a failure to display any morphological response to cold shock (NCU02636, peroxin-4), E) Thin: hyphal diameter narrows on cold shock (NCU03013, anchored cell wall protein-10), F) Dense: apical branching tighter than that normally displayed during cold shock (NCU07617, aconidiate-3), G) Cot-like: phenotype resembles that seen at the restrictive temperature of a temperature-sensitive colonial (cot) mutant strain. (NCU03901, peroxin-14), H) Weak: apical branching during cold shock which is less dense than normally observed (NCU01408, COP9 signalosome-3). Combinations of the above were sometimes observed as noted in Table 1.

Homeostasis in the face of temperature changes and more specifically the response to cold shock has been extensively studied in bacterial systems for over 20 years. DNA microarray transcription profiling of the cold shock response in *E. coli* by Phadtare and Inouye (2004) identified a collection of genes which alter their level of transcription in response to cold shock.

The hypothesis of this project was that the observed cold shock response of Neurospora is a consequence of a cellular response homologous to that induced by cold shock in bacteria. Under this hypothesis, the observed, transient morphological changes are a consequence of the fungal cell adjusting itself to growth in the cold via a manner which is shared in common with simpler organisms. This hypothesis was tested by screening Neurospora knockout strains impacting genes homologous to those identified in *E. coli* which alter their expression patterns in response to cold shock. In addition, a broader collection of selected knockout strains were screened to identify additional genes which play a role in the cold shock response and thus cold acclimatization. Together, the results of this screen provides the first molecular underpinning to the cold shock response in Neurospora,

## MATERIALS AND METHODS

### The Neurospora targeted deletion collection

As part of the Neurospora Genome Project, a collection of strains containing disruptions in presumptive genes was constructed (Colot *et al.* 2006). Strains representing deletions of most of the genes of the Neurospora genome are available from the Fungal Genetics Stock Center (McCluskey 2003). As each deletion strain has been altered in a single, previously identified, presumptive gene – going from phenotype to sequence is greatly simplified.

The accession numbers listed in Tables 1&2 represent the locus number of the gene subject to inactivation in the knockout strain under test. Every annotated gene in *Neurospora crassa* has been assigned a locus number of the form NCU#####. The gene identities reported in the tables are those associated with the genes as annotated on the FungiDB database as of July 2017: fungidb.org/fungidb/. The gene identities reported are based solely on the annotations currently associated with those strains and have not been independently confirmed by the authors of this study. Gene Ontologies reported are those determined by pantherdb.org (Mi *et al.* 2016) as of December 2017.

**Table 1:** Of 334 knockouts screened 115 were observed to alter the phenotype of the cold shock response. For each knockout strain tested (“ID”/NCU#####) we report the Cold Shock phenotype, the annotated gene function and gene abbreviation, the set of mutants the knockout came from (*E. coli* cold shock mutant ortholog, the Morphological or Hyphal growth plates from the FGSC, or the Transcription Factor plates from the FGSC) and the Gene Ontology categorizations for both Molecular Function and Biological Process.

| ID | CS Phenotype | Gene Function | Gene | Knockout set | GO: Molecular Function | GO: Biological Process |
| --- | --- | --- | --- | --- | --- | --- |
| NCU03938 | burst | alternative oxidase-5 | aod-5 | Morph/Hyph |  |  |
| NCU03070 | burst | hypothetical protein |  | Transc Factors | Binding | Biological Regulation |
| NCU01782 | burst | Ras guanyl-nucleotide exchange factor RasGEF |  | *E. coli* CS Orth |  | Cellular Process |
| NCU02133 | burst | superoxide dismutase-1 | sod-1 | Morph/Hyph | antioxidant/binding/catalytic Activity | Cellular Process/<br>Response to Stimulus |
| NCU01213 | burst | superoxide dismutase-2 | sod-2 | *E. coli* CS Orth | Binding | Developmental Process |
| NCU03623 | burst | ubiquitin-conjugating enzyme E |  | Morph/Hyph | Binding | Cellular Process/<br>Metabolic Process |
| NCU04242 | burst/dense | period-6 | prd-6 | Morph/Hyph | Binding/Catalytic Activity | Biological Regulation/<br>Cellular Process/<br>Metabolic Process |
| NCU07728 | burst/thin | siderophore regulation | sre | Morph/Hyph |  |  |
| NCU03901 | cot-like | peroxin 14 | pex14 | Morph/Hyph | Binding | Cellular Component<br>Organization or<br>Biogenesis/ Cellular<br>Process/ Localization/<br>Metabolic Process |
| NCU07617 | dense | aconidiate-3 | acon-3 | Morph/Hyph |  | Biological Regulation/<br>Developmental Process/<br>Reproduction |
| NCU05410 | dense | arginine-5 | arg-5 | *E. coli* CS Orth | Binding | Cellular Process/<br>Metabolic Process |
| NCU02114 | dense | G1/S-specific cyclin Cln1 |  | Morph/Hyph | Binding | Cellular Process |
| NCU00144 | dense | hypothetical protein |  | Transc Factors |  |  |
| NCU03120 | dense | hypothetical protein |  | Transc Factors |  |  |
| NCU03356 | dense | hypothetical protein |  | Transc Factors |  |  |
| NCU03417 | dense | hypothetical protein |  | Transc Factors |  |  |
| NCU03905 | dense | hypothetical protein |  | Transc Factors |  |  |
| NCU03962 | dense | hypothetical protein |  | Transc Factors | Binding | Cellular Process/<br>Metabolic Process |
| NCU06990 | dense | hypothetical protein |  | Transc Factors |  |  |
| NCU01154 | dense | submerged protoperithecia-1 | sub-1 | Transc Factors |  |  |
| NCU04899 | dense | tricarboxylic acid-15 | tca-15 | *E. coli* CS Orth | Catalytic Activity | Metabolic Process |
| NCU03415 | fail | aldehyde dehydrogenase | CBS-3 | *E. coli* CS Orth |  |  |
| NCU11289 | fail | aldo-keto reductase |  | *E. coli* CS Orth | Catalytic<br>Activity/transporter Activity |  |
| NCU00097 | fail | BEAK-1 | bek-1 | Transc Factors |  |  |
| NCU02017 | fail | CBF/NF-Y family transcription<br>factor | ada-2 | Transc Factors |  |  |
| NCU00056 | fail | condensing enzyme with<br>mitochondrial function | cem-1 | *E. coli* CS Orth |  |  |
| NCU00467 | fail | COP9 signalosome-5 | csn-5 | Morph/Hyph | Binding | Metabolic Process |
| NCU06068 | fail | fungal specific transcription<br>factor | col-25 | Transc Factors |  |  |
| NCU07788 | fail | fungal specific transcription<br>factor | col-26 | Transc Factors |  |  |
| NCU07945 | fail | fungal specific transcription<br>factor | tah-4 | Transc Factors |  |  |
| NCU07947 | fail | glycolipid transfer protein HET-C2 |  | Morph/Hyph |  | Localization/ Metabolic<br>Process |
| NCU05927 | fail | GTP-binding protein GUF1 | GTP-7 | *E. coli* CS Orth |  |  |
| NCU00528 | fail | hyphal anastomosis-4 | ham-4 | *E. coli* CS Orth |  |  |
| NCU07561 | fail | hypothetical protein |  | Transc Factors |  |  |
| NCU09120 | fail | lysine-specific histone<br>demethylase Aof2 |  | *E. coli* CS Orth | Binding | Cellular Process/<br>Metabolic Process |
| NCU09830 | fail | menadione-induced gene-12 | mig-12 | Morph/Hyph | Catalytic Activity | Cellular Process/<br>Metabolic Process |
| NCU09842 | fail | mitogen activated protein kinase-1 | mak-1 | Morph/Hyph | Catalytic Activity/signal<br>transducer activity | Biological Regulation/<br>Cellular Process/<br>Response to Stimulus/<br>Metabolic Process |
| NCU03314 | fail | mob2-like-a | mob-2a | Morph/Hyph | Binding/Catalytic Activity | Cellular Process |
| NCU09975 | fail | multidrug resistance protein 3 |  | *E. coli* CS Orth |  |  |
| NCU08294 | fail | nitrogen assimilation transcription factor nit-4 | nit-4 | Transc Factors |  |  |
| NCU03277 | fail | peroxin 10 | pex10 | Morph/Hyph |  | Cellular Component Organization or Biogenesis/ Cellular Process/ Localization |
| NCU02636 | fail | peroxin 4 | pex4 | Morph/Hyph |  |  |
| NCU01004 | fail | phosphatidylserine decarboxylase proenzyme | CHOL-15 | *E. coli* CS Orth | Catalytic Activity | Cellular Process/ Metabolic Process |
| NCU07832 | fail | pre-mRNA processing splicing factor 8 | msp-39 | *E. coli* CS Orth | Catalytic Activity | Cellular Component Organization or Biogenesis/ Cellular Process/ Metabolic Process |
| NCU06028 | fail | quinic acid utilization activator | qa-1F | Transc Factors |  |  |
| NCU06205 | fail | regulator of conidiation-1 | rco-1 | Morph/Hyph |  |  |
| NCU06145 | fail | RING-6 | RING-6 | Morph/Hyph |  |  |
| NCU02214 | fail | TAH-2 | tah-2 | Transc Factors |  |  |
| NCU10008 | fail | tricarboxylic acid-14 | tca-14 | *E. coli* CS Orth |  | Cellular Process/ Metabolic Process |
| NCU02356 | fail | white collar 1 | wc-1 | Transc Factors | Binding |  |
| NCU02173 | fail | zinc finger transcription factor-52 | znf-52 | Transc Factors |  |  |
| NCU05591 | thin | ABC transporter CDR4 |  | Morph/Hyph | Catalytic Activity/transporter Activity | Cellular Process/ Metabolic Process |
| NCU03013 | thin | anchored cell wall protein-10 | acw-10 | Morph/Hyph | antioxidant/binding/catalytic Activity | Cellular Process/ Response to Stimulus |
| NCU02333 | thin | arginase-1 | aga-1 | Morph/Hyph | Binding/Catalytic Activity | Cellular Process/ Metabolic Process |
| NCU03184 | thin | C2H2 conidiation transcription factor FlbC |  | Morph/Hyph |  |  |
| NCU07075 | thin | calcium exchanger | cax | Morph/Hyph | transporter activity | Biological Regulation/<br>Cellular Process |
| NCU05770 | thin | catalase-2 | cat-2 | *E. coli* CS Orth | antioxidant/binding | Cellular Process/<br>Response to Stimulus/<br>Metabolic Process |
| NCU05051 | thin | COL-23 | col-23 | Transc Factors |  |  |
| NCU00830 | thin | ctr copper transporter | tcu-1 | Morph/Hyph |  |  |
| NCU08216 | thin | cystathionine beta-synthase | MET-11 | *E. coli* CS Orth | Binding | Cellular Process/<br>Metabolic Process |
| NCU03076 | thin | delta-1-pyrroline-5-carboxylate<br>dehydrogenase |  | Morph/Hyph |  |  |
| NCU08968 | thin | dimethyladenosine transferase |  | *E. coli* CS Orth |  | Cellular Component<br>Organization or<br>Biogenesis/ Cellular<br>Process/ Metabolic<br>Process |
| NCU01772 | thin | DNA-directed RNA polymerase<br>III polypeptide | rpo-10 | *E. coli* CS Orth |  | Cellular Process/<br>Metabolic Process |
| NCU02542 | thin | embden-meyerhof pathway-1 | emp-1 | Morph/Hyph | Catalytic Activity | Biological Regulation/<br>Cellular Process/<br>Metabolic Process |
| NCU01744 | thin | enhancer-2 of am | en(am)-2 | *E. coli* CS Orth | Catalytic Activity | Cellular Process/<br>Metabolic Process |
| NCU04264 | thin | extracellular developmental<br>signal biosynthesis protein FluG |  | Morph/Hyph | Binding | Cellular Process/<br>Metabolic Process |
| NCU04140 | thin | FK506 resistant-2 | fkr-2 | *E. coli* CS Orth | Binding/Catalytic Activity | Cellular Process/<br>Metabolic Process |
| NCU09930 | thin | folic acid synthesis protein | fol-9 | *E. coli* CS Orth | transporter activity | Cellular Process/<br>Metabolic Process |
| NCU05606 | thin | glucosidase 2 subunit beta | GHX-4 | *E. coli* CS Orth | Binding/Catalytic Activity | Biological Regulation/<br>Metabolic Process |
| NCU01528 | thin | glyceraldehyde-3-phosphate<br>dehydrogenase-1 | gpd-1 | *E. coli* CS Orth | Catalytic Activity | Metabolic Process |
| NCU06005 | thin | glycerol kinase | GLK-1 | *E. coli* CS Orth |  |  |
| NCU02630 | thin | heat shock protein 78 | hsp78 | *E. coli* CS Orth |  |  |
| NCU07156 | thin | histidine-6 | his-6 | *E. coli* CS Orth |  |  |
| NCU02556 | thin | histone acetyl transferase-2 | hat-2 | *E. coli* CS Orth |  | Cellular Component Organization or Biogenesis/ Cellular Process/ Metabolic Process |
| NCU01629 | thin | hypothetical protein |  | Transc Factors |  |  |
| NCU04669 | thin | hypothetical protein homologous to Bactericidal permeability-increasing protein |  | Morph/Hyph |  |  |
| NCU04561 | thin | melanization defective-1 | mld-1 | Transc Factors |  |  |
| NCU09767 | thin | membrane transporter |  | *E. coli* CS Orth |  |  |
| NCU04791 | thin | menadione-induced gene-10 | mig-10 | *E. coli* CS Orth |  |  |
| NCU05151 | thin | phosphoketolase | PHK-1 | *E. coli* CS Orth |  |  |
| NCU06342 | thin | phospholipase D | PLA-5 | *E. coli* CS Orth |  | Cellular Process/ Localization/ Locomotion |
| NCU05295 | thin | proteasome catalytic alpha-5 | pca-5 | Morph/Hyph |  | Cellular Process/ Metabolic Process |
| NCU09366 | thin | proteasome catalytic beta-6 | pcb-6 | Morph/Hyph | Catalytic Activity | Cellular Process/ Metabolic Process |
| NCU01613 | thin | protoperithecium-2 | pp-2 | Morph/Hyph | Catalytic Activity | Biological Regulation/ Cellular Component Organization or Biogenesis/ Cellular Process/ Metabolic Process |
| NCU02260 | thin | regulatory particle, ATPase-like-3 | rpt-3 | Morph/Hyph |  |  |
| NCU02055 | thin | uridine nucleosidase Urh1 | NUS-1 | *E. coli* CS Orth |  |  |
| NCU07705 | thin/fail | C6 finger domain-containing protein | clr-1 | Transc Factors |  |  |
| NCU08000 | thin/fail | cutinase transcription factor 1 alpha | far1 | Transc Factors |  |  |
| NCU05536 | thin/fail | hypothetical protein |  | Transc Factors | Binding/Catalytic Activity | Cellular Component Organization or Biogenesis/ Cellular Process/ Response to Stimulus/ Metabolic Process |
| NCU08651 | thin/fail | zinc binuclear cluster-type protein | col-27 | Transc Factors |  |  |
| NCU07732 | thin/fail | arginine-2 | arg-2 | *E. coli* CS Orth | Binding/Catalytic Activity | Cellular Process/<br>Metabolic Process |
| NCU04117 | thin/fail | ATP-dependent permease MDL2 | ABC-7 | *E. coli* CS Orth |  |  |
| NCU06659 | thin/fail | GTP-binding protein | GTP-3 | *E. coli* CS Orth |  | Cellular Process/<br>Metabolic Process |
| NCU08693 | thin/fail | heat shock protein 70-5 | hsp70-5 | *E. coli* CS Orth | Binding | Cellular Process/<br>Response to Stimulus/<br>Metabolic Process |
| NCU10760 | thin/fail | jumonji domain-containing protein 5 |  | *E. coli* CS Orth |  |  |
| NCU08858 | thin/fail | MFS alpha-glucoside transporter | SUT-1 | *E. coli* CS Orth | transporter activity | Cellular Process/<br>Localization |
| NCU00793 | thin/fail | trehalose phosphate synthase | GT20-2 | *E. coli* CS Orth | Catalytic Activity | Cellular Process/<br>Response to<br>Stimulus/Metabolic<br>Process |
| NCU08336 | thin/fail | tricarboxylic acid-12 | tca-12 | *E. coli* CS Orth | Catalytic Activity | Cellular Process/<br>Localization/ Metabolic<br>Process |
| NCU00771 | thin/fail | UBX domain-containing protein 7 |  | *E. coli* CS Orth |  |  |
| NCU04583 | weak | acetyltransferase |  | *E. coli* CS Orth | Catalytic Activity |  |
| NCU00499 | weak | all development altered-1 | ada-1 | Transc Factors |  |  |
| NCU00567 | weak | arginine-6 | arg-6 | *E. coli* CS Orth | Binding | Cellular Process/<br>Metabolic Process |
| NCU04303 | weak | asparagine synthetase 2 | asn-1 | *E. coli* CS Orth | Binding/Catalytic Activity | Cellular Process/<br>Metabolic Process |
| NCU00919 | weak | ATP-dependent RNA helicase rok-1 | drh-16 | *E. coli* CS Orth |  |  |
| NCU08933 | weak | cellular nucleic acid-binding protein |  | *E. coli* CS Orth |  |  |
| NCU01408 | weak | COP9 signalosome-3 | csn-3 | Morph/Hyph |  | Cellular Process/<br>Metabolic Process |
| NCU01625 | weak | DNA repair helicase RAD3 | DNR-10 | *E. coli* CS Orth | Binding/Catalytic Activity | Cellular Process/<br>Metabolic Process |
| NCU07027 | weak | glycogen phosphorylase | GYP-1 | *E. coli* CS Orth | Binding/Catalytic Activity | Cellular Process/<br>Metabolic Process |
| NCU06523 | weak | glycosylhydrolase family 13-4 | gh13-4 | *E. coli* CS Orth | Catalytic Activity | Metabolic Process |
| NCU01589 | weak | heat shock protein 60 | hsp60 | *E. coli* CS Orth | Binding | Cellular Component<br>Organization or<br>Biogenesis/ Cellular<br>Process/ Metabolic<br>Process |
| NCU05909 | weak | hypothetical protein |  | Transc Factors |  |  |
| NCU08439 | weak | leptomycin B resistance protein<br>pmd1 | ABC-2 | *E. coli* CS Orth | transporter activity | Cellular Process/<br>Metabolic Process |
| NCU00565 | weak | lipoic acid synthetase | LIA-1 | *E. coli* CS Orth | Catalytic Activity | Cellular Process/<br>Metabolic Process |
| NCU04339 | weak | ribokinase | RIK-8 | *E. coli* CS Orth | Catalytic Activity | Cellular Process/<br>Metabolic Process |
| NCU03894 | weak | serine/threonine protein kinase-4 | stk-4 | Morph/Hyph | Binding/signal transducer<br>activity | Biological Regulation/<br>Cellular Process/<br>Developmental Process/<br>Response to Stimulus |
| NCU06017 | weak | thiosulfate sulfurtransferase | TST-1 | *E. coli* CS Orth |  | Localization/ Response<br>to Stimulus/ Metabolic<br>Process |
| NCU10053 | weak | thymidylate synthase | pyr-8 | *E. coli* CS Orth |  |  |
| NCU08658 | weak | zinc finger transcription factor-50 | znf-50 | Transc Factors |  |  |

**Table 2:**
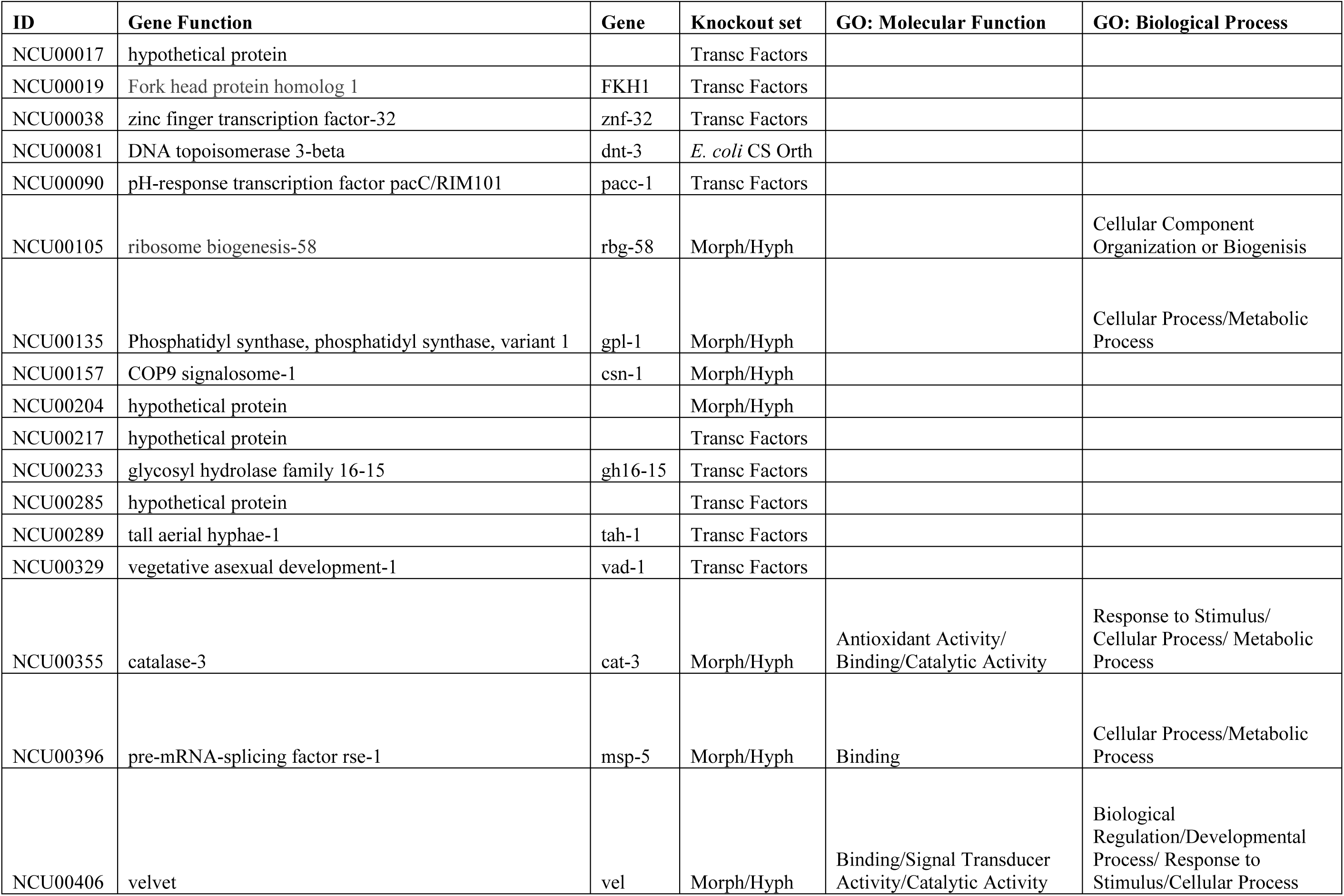
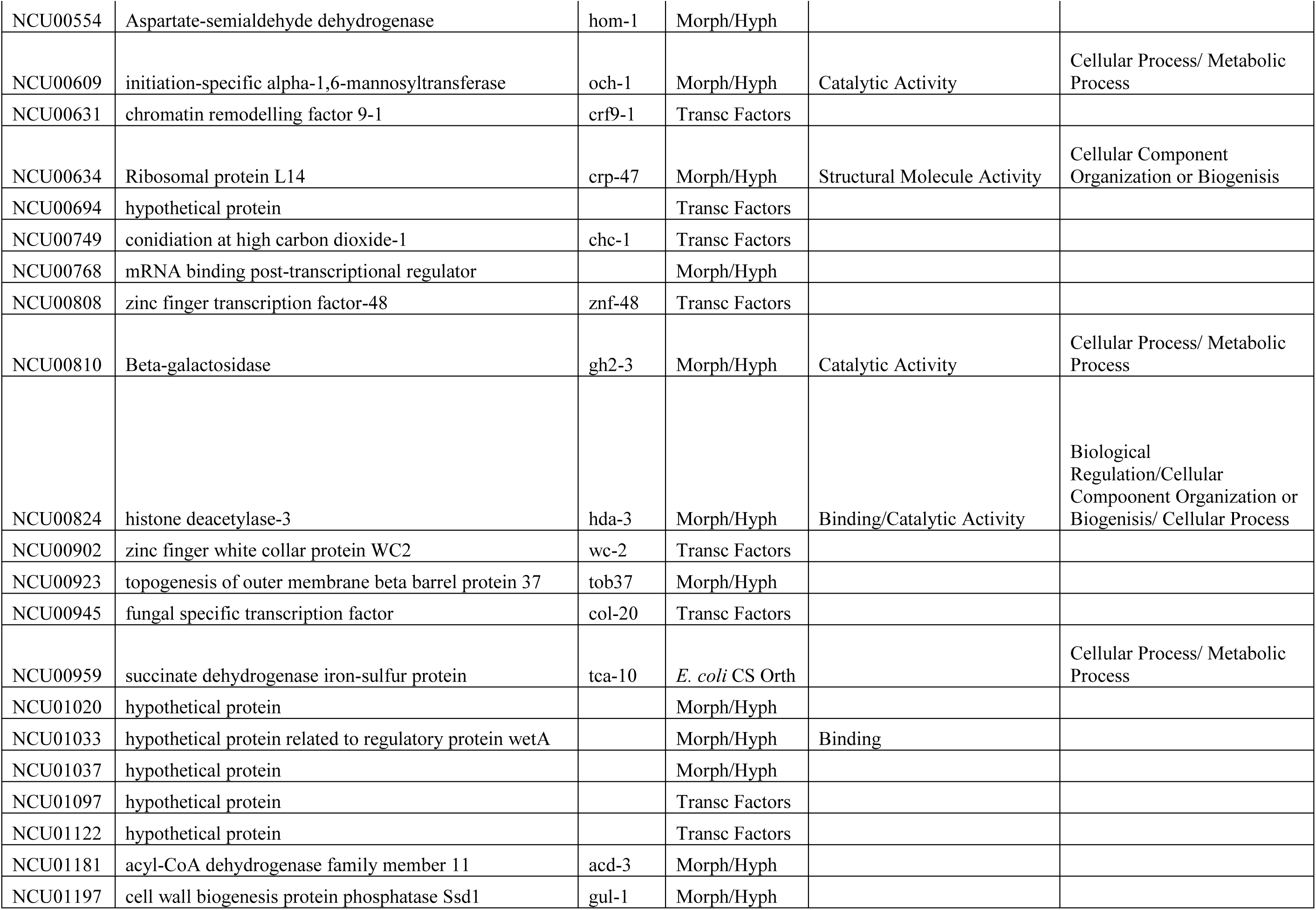
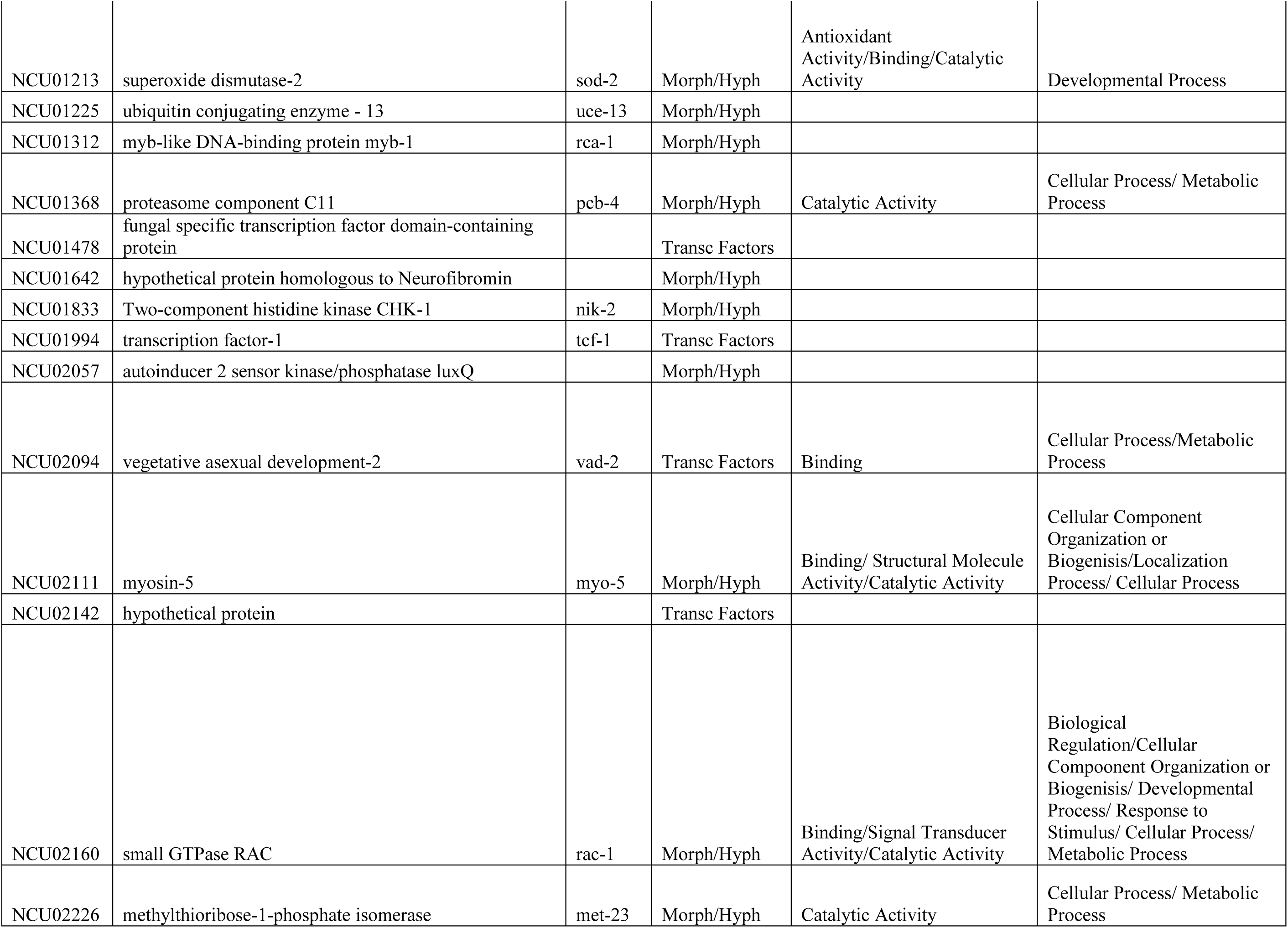
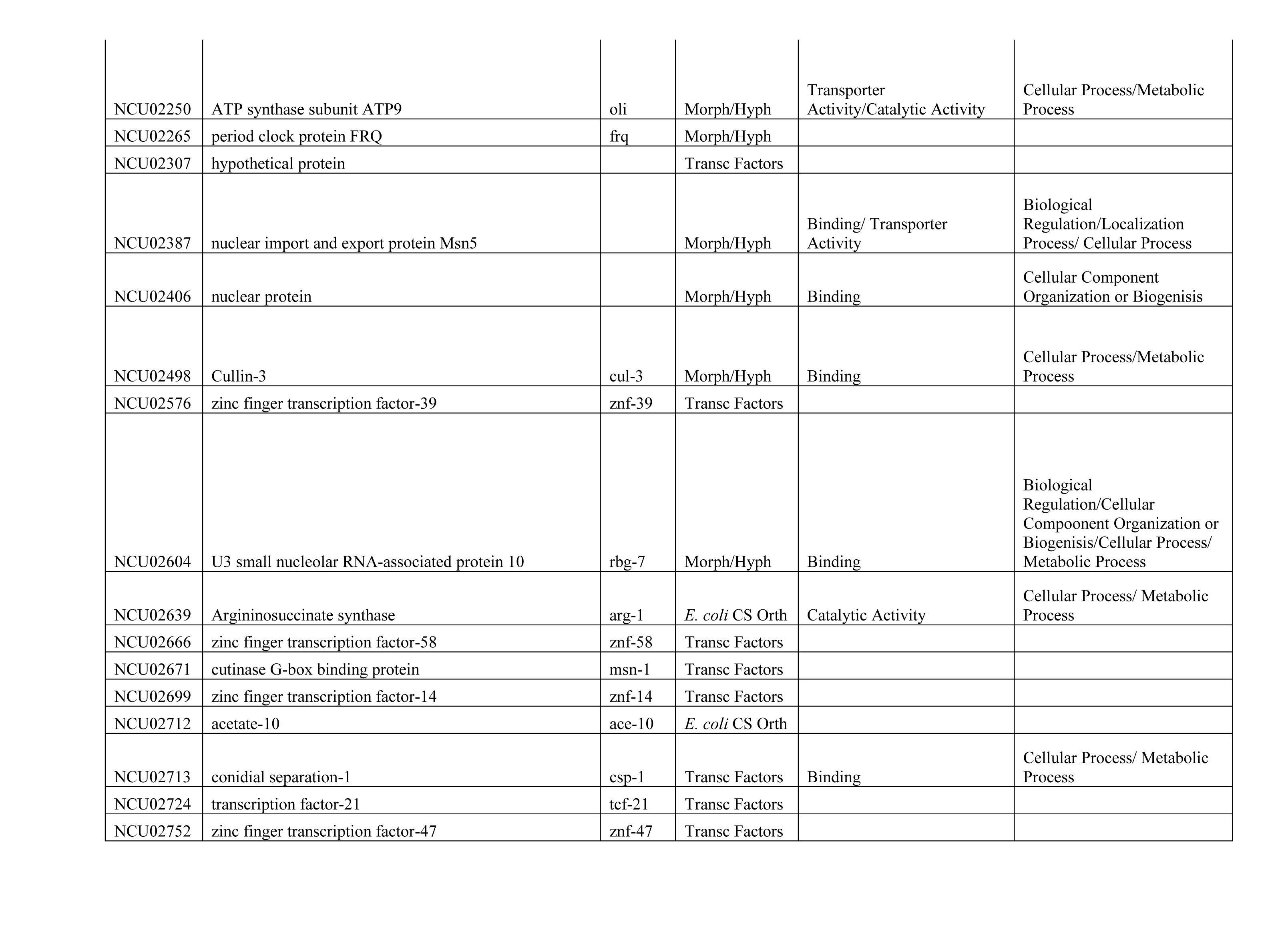
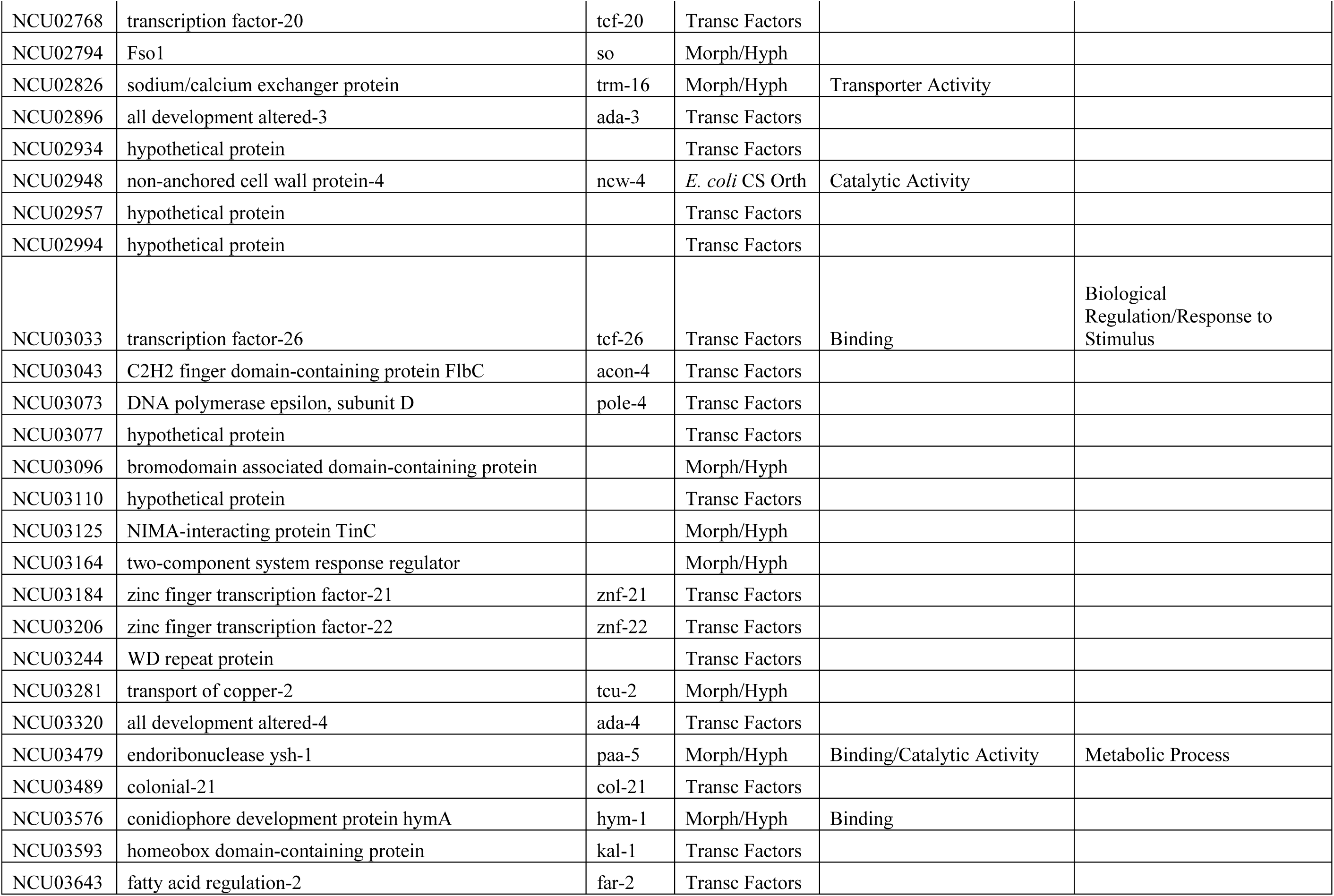
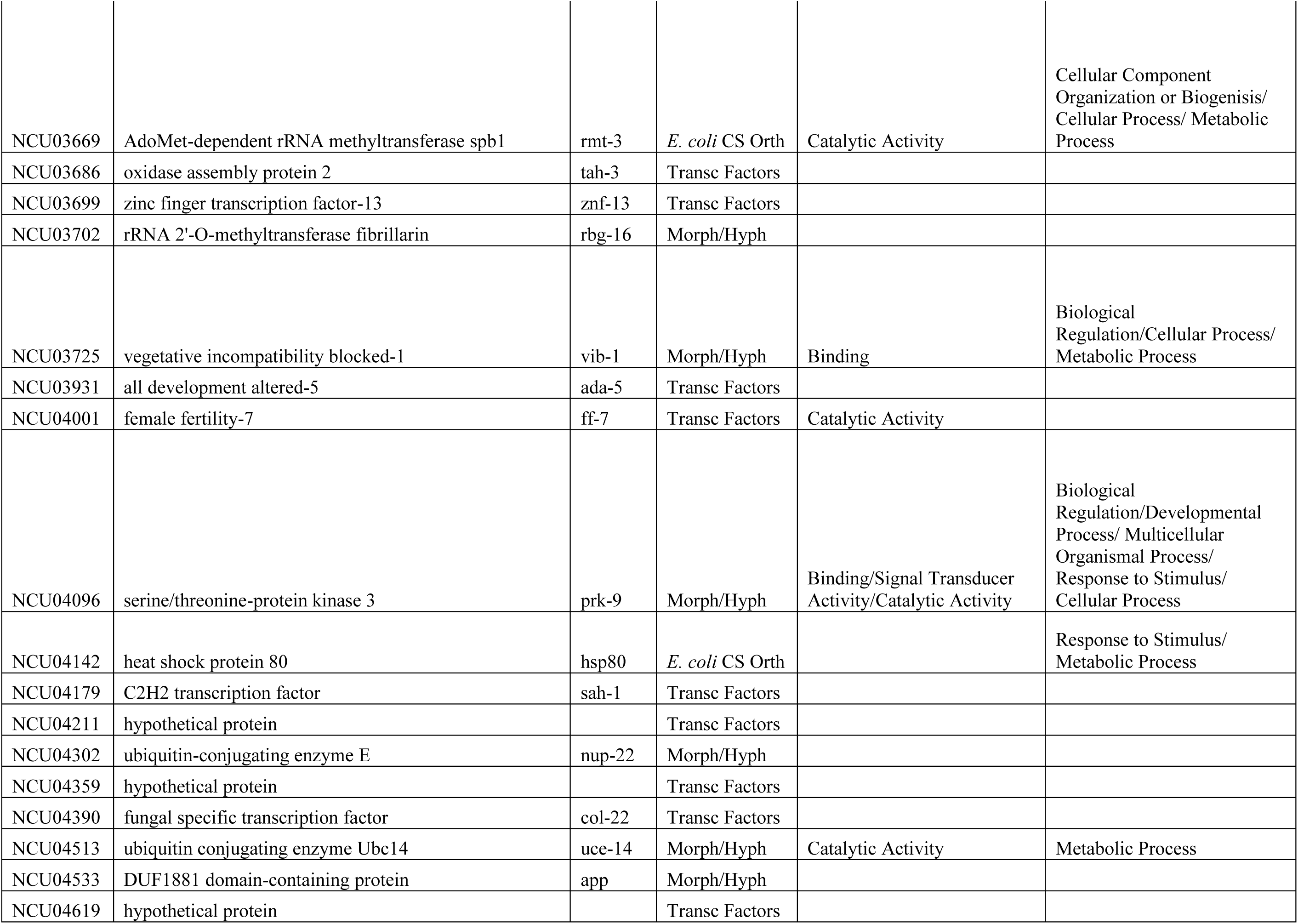
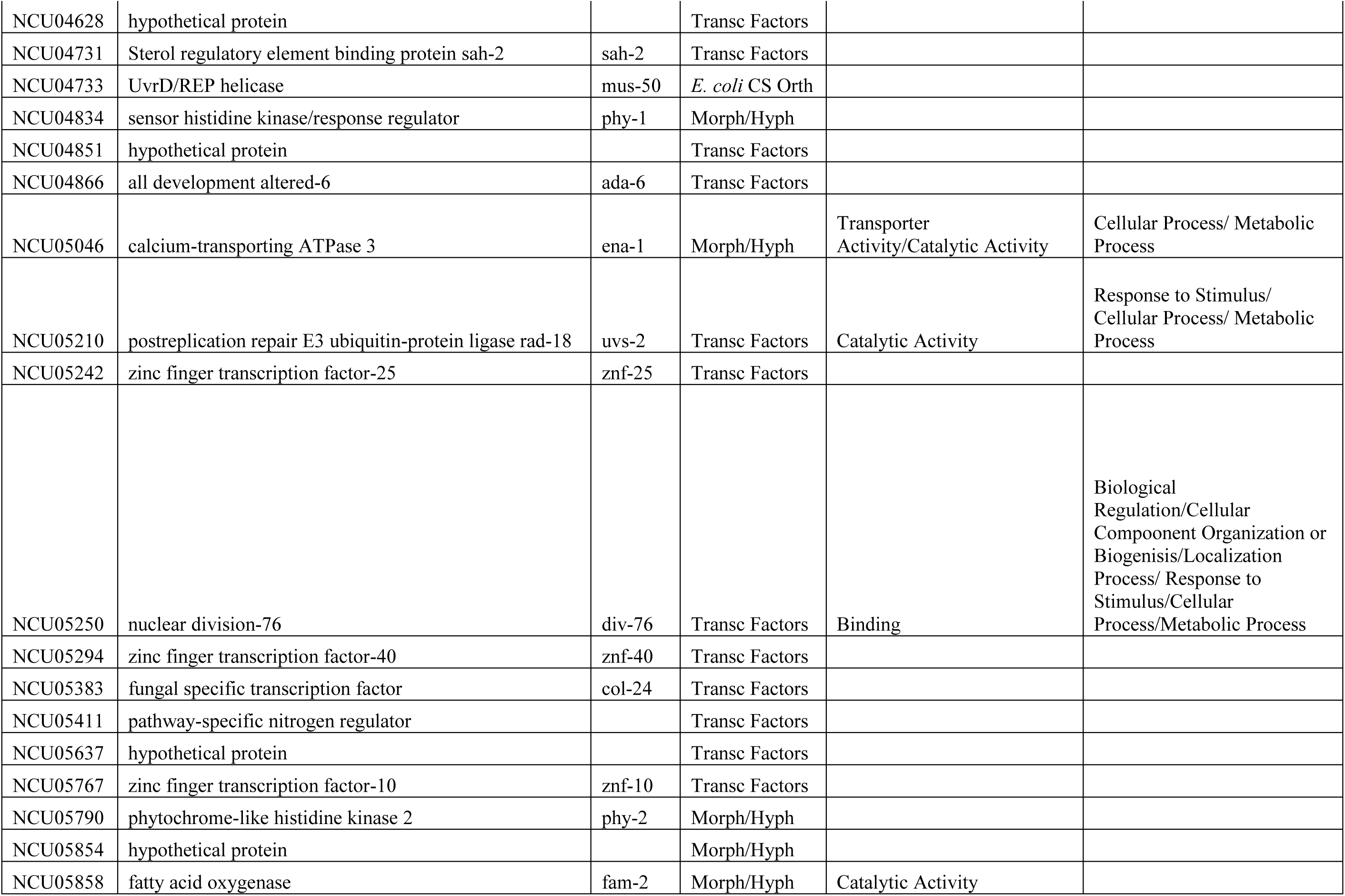
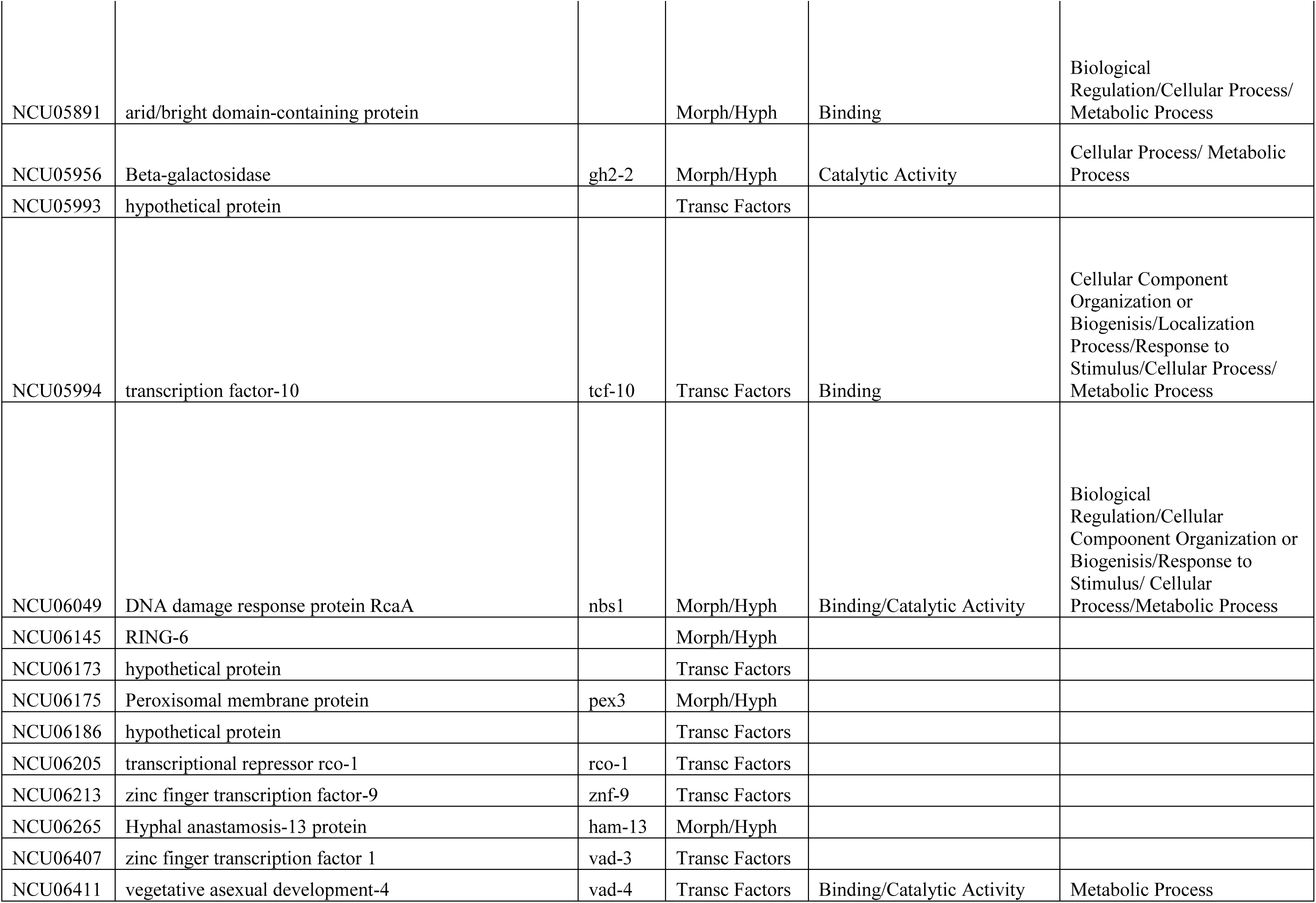
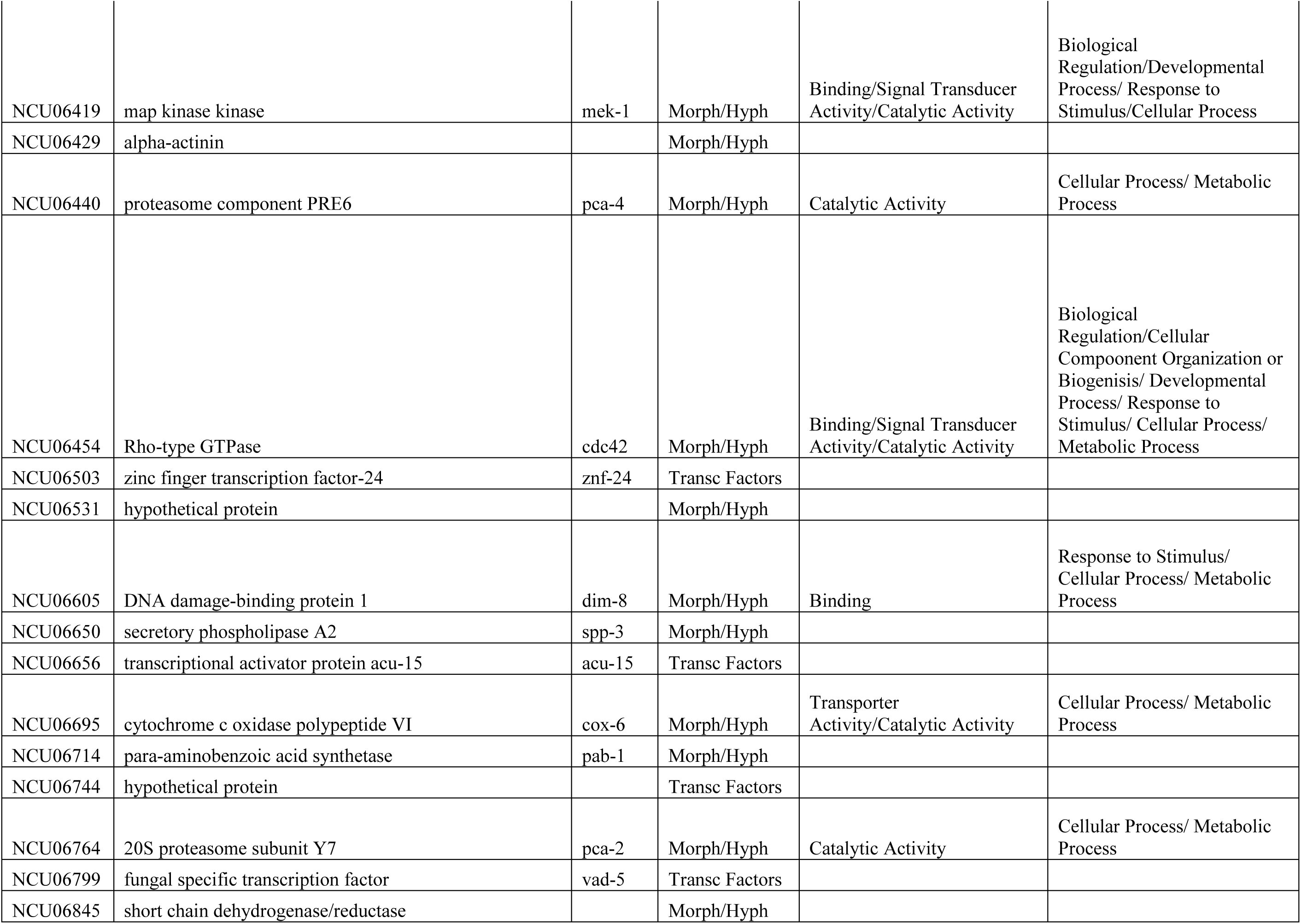
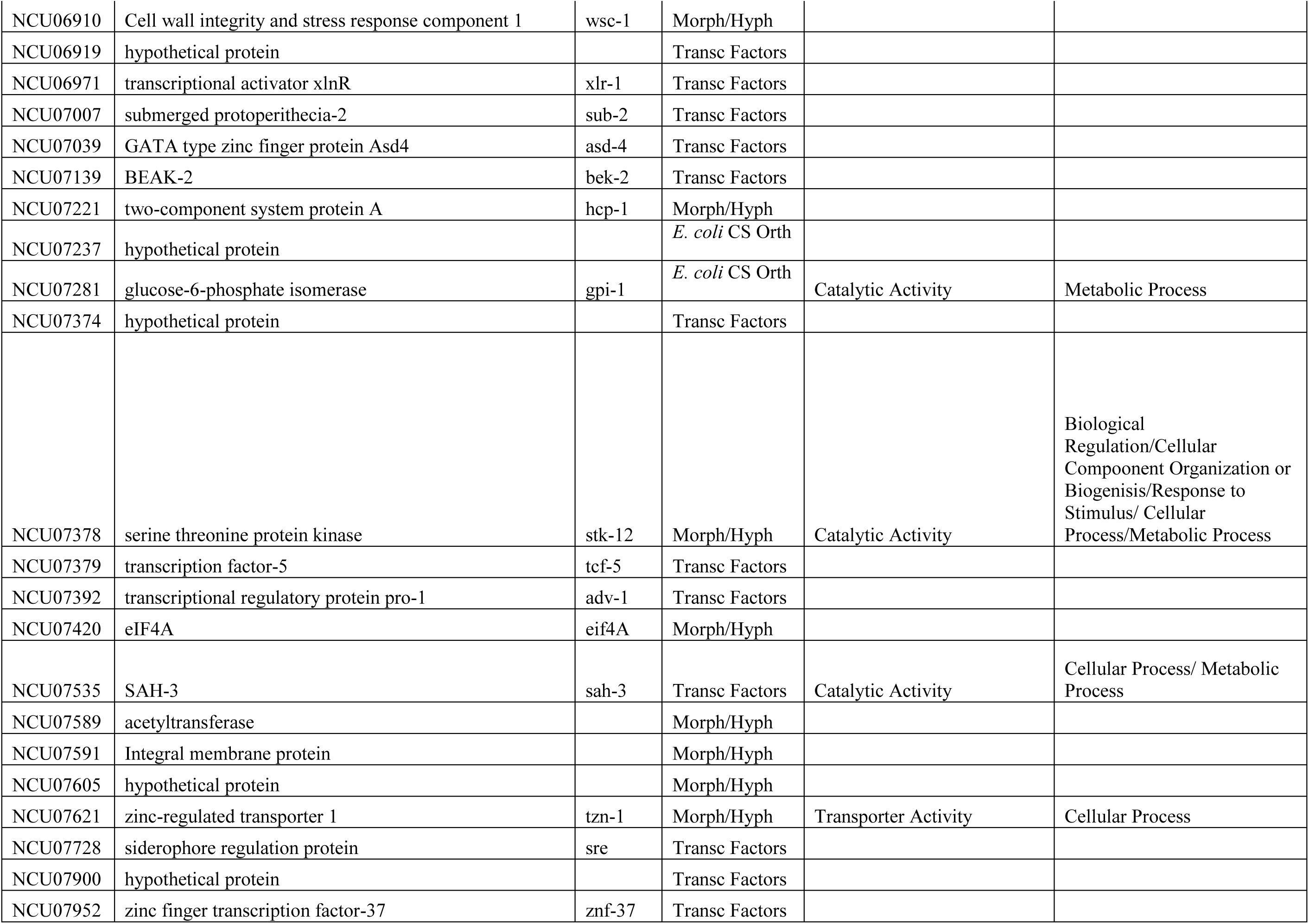
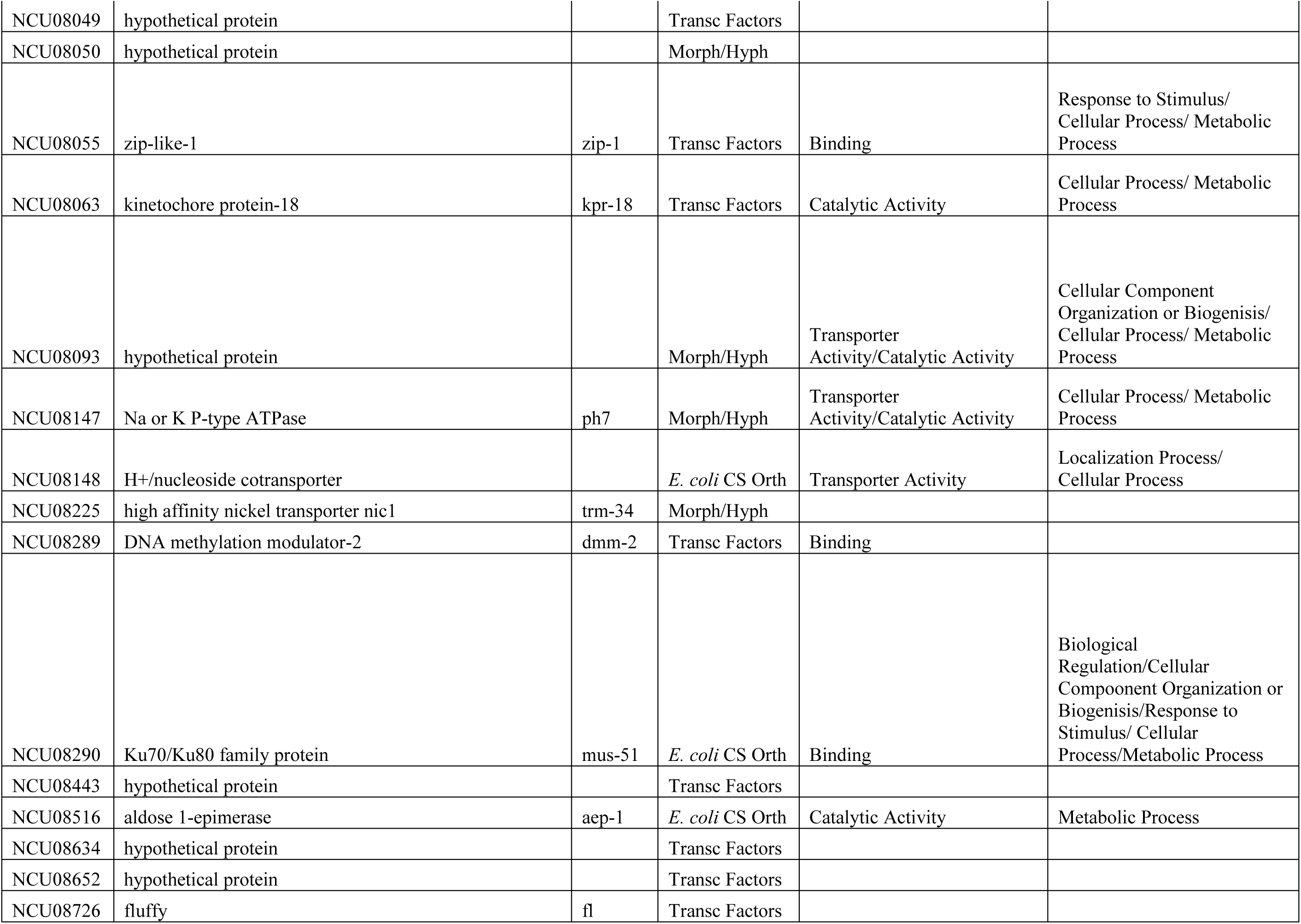
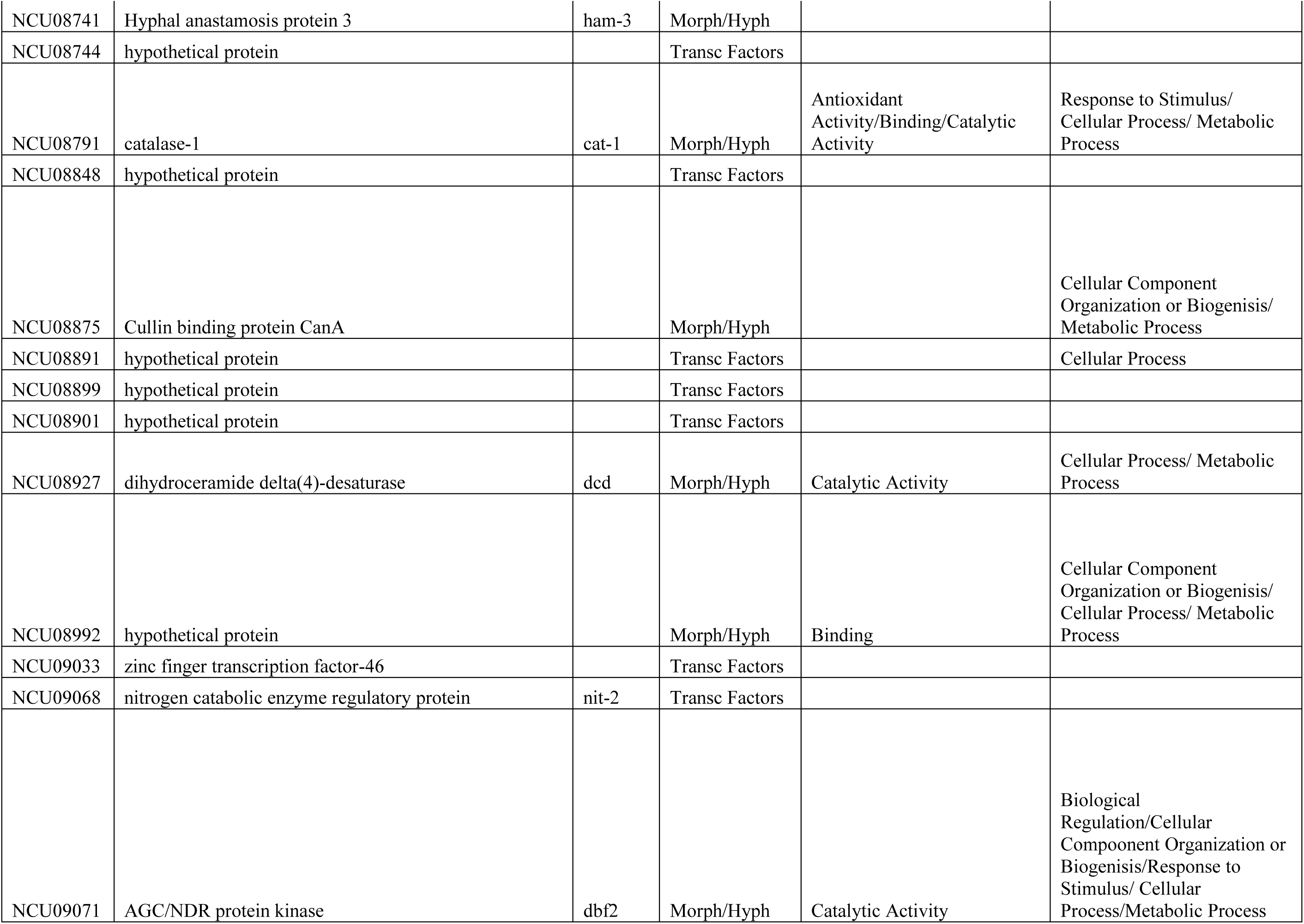
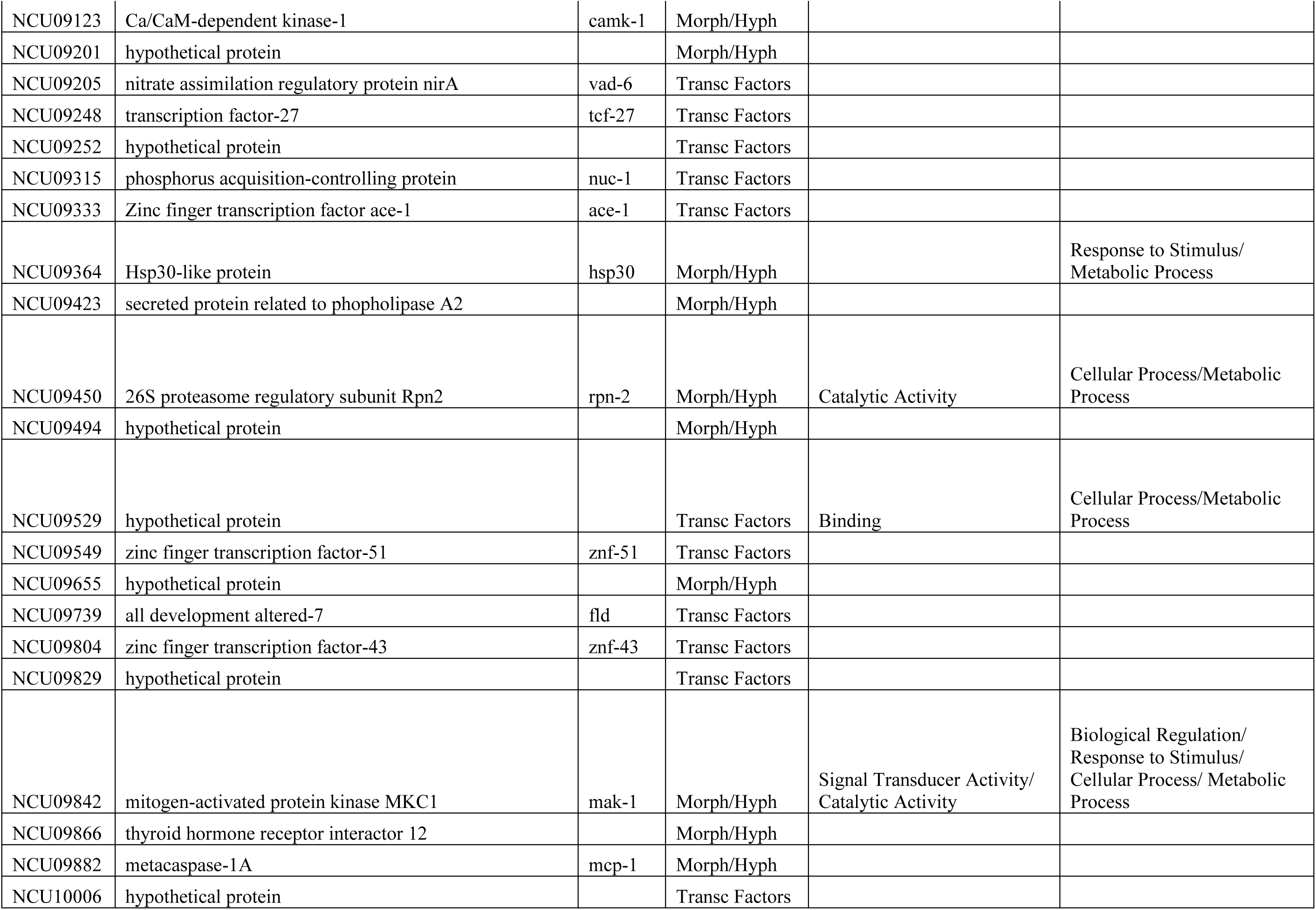
Of knockouts screened, 229 presented no change to the cold shock morphology. Columns are the same as for Table 1.

### Knockout sets selected to be subjected to screen

A screen of the entire library was determined to be impractical. We instead screened an abbreviated subsection of the library chosen to be more likely to yield positive responses. These fall into three basic sets.

The first set are knockouts of genes homologous to those which show altered transcription in *E. coli* when subjected to cold shock (Phadtare and Inouye 2004). The protein sequences of *E. coli* genes identified by were retrieved from the *E. coli* database (ecocyc.org/). These amino acid sequences were then fed into a BLAST search on the NIH NCBI site (blast.ncbi.nlm.nih.gov/Blast.cgi) with the output limited to Neurospora sequences in order to identify their nearest Neurospora homologs. These homologs were then searched on FungiDB to determine which had knockout strains available. From this final list, 68 were selected for screening in this study. This set was selected to determine the degree of relationship between the cold shock response in *E. coli* and Neurospora.

Second, two previously organized sets of knockouts generally associated with hyphal growth and morphology and available from the FGSC were included in this screen. One set (identified as “plate 29 – morphologicals” by the FGSC) contained strains with knockouts known to cause morphological changes. The second set (identified as “Hyphal Growth Set” by the FGSC) contained strains with knockouts in genes homologous to genes in yeast known to affect polar growth. A total of 129 strains from these two sets were screened.

The last set consists of knockouts of known transcription factors in Neurospora. This collection is available as a set from the Fungal Genetics Stock Center (McCluskey 2003). It was selected for this screen to determine which transcription factors play a role in signaling to the cell that cold adaptation genes must be activated. A total of 147 strains from this set were screened.

### Media

Media and culturing procedures were those described in Davis & deSerres (1970). Growth described as being on “minimal” was in plates containing Vogel’s minimal medium (Davis & deSerres 1970) with 2% agar.

### Screen

The selected knockout strains were subjected to a screen looking for altered responses to cold shock. Wild-type Neurospora progresses through a three-stage response following a shift into the cold. To induce the cold shock response, we initially grew strains at 33°C and shifted to 4°C. We selected 33°C as our “normal” temperature as the cold shock response has previously been demonstrated to be dependent on the degree of the temperature shift the hypha are subjected to (Watters et al. 2000). The larger temperature shift used here would be expected to result in tighter branching during the apical phase. We decided this was desirable as it would make any variations from the normal cold shock response more visible and easier to identify in the screen. Strains were inoculated by dropping a suspension of conidia onto Vogel’s Minimal Medium and incubated overnight at 33°C. The next morning plates were moved to 4°C. After an overnight incubation at 4°C, the strain’s response to cold shock was photographed and evaluated. Variations in the cold shock response from that of wild-type Neurospora were judged qualitatively. Knockouts were subjected to cold shock and photographed a minimum of three independent runs on separate days to assure consistency of the response within a strain.

### Photomicroscopy

Growing cultures were examined and photographed using a Motic 10MP digital camera attached to a Wolfe Beta Elite trinocular microscope. Photographs were taken of well separated, leading hyphae. All photomicrographs were taken using 40x magnification.

### Phenotypes scored

The morphology of strains following cold shock was scored visually by comparing collections of photographs of cold shock in a given strain to the response seen with a wild type strain (*Neurospora crassa* Oak Ridge). Those with altered responses were then further categorized visually into the groups reported in Table 1 “CS phenotype.”

## RESULTS AND DISCUSSION

When Neurospora is subjected to a rapid temperature downshift, a 3-phase response is observed (Figure 1, Watters *et al.* 2000, Watters 2013). The initial response to cold shock is the growth of a single longer than normal unbranched segment. This was termed the “Lag” phase of the response. This phase is followed by a series of closely spaced apical branch points, termed the “Apical” phase. Apical branch formation has been previously associated with the disruption and attempted reorganization of the normal tip-growth apparatus (Reynaga-Peña *et al.* 1995, Riquelme & Bartnicki-Garcia 2004), a mechanism distinct from that thought to be involved in lateral branching. Finally, with continued incubation at the lower temperature, the colony returns to lateral branching, termed the “Recovery” phase. Growth in this phase of the response resembles that which would be seen had the colony been grown at 4°C (or any other fixed temperature) continuously (Watters et al 2000). Thus, the cold shock response appears to be a temporary disturbance to a homeostatic system which maintains branch density at a constant, evolutionarily favored, value. The morphological effects of cold shock are the indirect consequence of this system’s staged process of adjusting cellular conditions in order to compensate for the new growth temperature.

### Cold shock response of *E. coli*

Homeostasis in the face of temperature changes and more specifically the response to cold shock has been extensively studied in bacterial systems for over 20 years. The effect of cold shock is manifest in multiple cellular systems including: membrane rigidity (Shivaji & Prakash 2010), stability of secondary structures in DNA/RNA (Phadtare 2004), efficiency of protein folding (Phadtare 2004) and ribosome function (Gualerzi *et al.* 2011). While much remains to be described in these systems, cold shock appears to result in a multi-stage response (Phadtare 2004). First, a lag period in which growth and translation of proteins generally cease. This is followed by an adjustment phase in which specific cold-shock proteins which compensate for the changes brought on by the cold are preferentially translated (Giuliodori *et al.* 2004). In the final stage, growth continues otherwise normally, but at a reduced rate. DNA microarray transcription profiling of the cold shock response in *E. coli* by Phadtare and Inouye (2004) has shown that several hundred genes respond to cold shock, either being transiently induced/repressed or showing prolonged induction/repression. Analogous responses to cold shock and/or cold acclimation have been observed in diverse organisms including plants (Guy 1999) and animals (Canclini & Esteves 2007). Attempts to uncover cold shock proteins in fungi (Fang & Leger 2010) have met with mixed success.

### Connections between cold shock in bacteria and Neurospora

It is tempting to draw parallels between what is known about cold shock in bacterial systems and the observed response of Neurospora to similar cold shocks. Many of the systems affected during bacterial cold shock would be expected to impact fungal tip growth and branching (e.g. membrane fluidity). In addition, the nature and timing of the two responses are similar. Both can be adjusted by changing the intensity of the cold shock with more mild shocks (lower temperature differences) producing more mild responses and more severe shocks (larger temperature differences) producing more severe responses. Furthermore, the dynamics of the responses parallel each other. In each, there is a multistage response. There is an initial response which is transient in nature, followed by a more long-term response which largely represents a return to normal growth.

During the initial study of the cold shock response in Neurospora (Watters et al 2000), it was observed that two classical morphological mutants (most notably “granular” and “delicate”) produced altered responses to cold shock (not reported), demonstrating that mutants could be obtained which influenced this process. We chose to screen mutants from the Neurospora knockout library for their cold shock response in order to provide a genetic grounding to this process which has, thus far, been lacking. We chose to use the mutants of the knockout library instead of the products of a random mutagenesis as the knockouts allow an immediate identification of gene function in most cases.

Knockout strains displaying an altered morphological response to cold shock were classified according to the specific variation they displayed. Examples are shown in Figure 1. The “burst” phenotype was defined as displaying a large number of growing tips which stop growing, swell and then structurally fail leaving a pool of cytoplasm at the tip. The “fail” phenotype was defined as failing to display the apical branch phase characteristic of cold shock. In the “fail” response, growth proceeds normally with lateral branching following cold shock. The “thin” phenotype was defined by a very rapid decrease in hyphal diameter following cold shock. It was common to observe “thin” in combination with other altered cold shock responses. The “dense” phenotype was defined by displaying apical branching with visibly shorter distances between branch points following cold shock relative to the response in wild-type. The “weak” phenotype was defined as the opposite – an apical branch phase with visibly longer distances between branch points relative to wild-type following cold shock. Finally, the “cot-like” phenotype was characterized by a lack of apical branching, but a shift to tightly spaced lateral branches which morphologically resembled the growth of the traditional *cot* mutants at the restrictive temperature.

### Screen of *E. coli* cold shock gene homolog knockout set

A total of 68 Neurospora strains with knockouts of genes homologous to *E. coli* genes which alter transcription in response to cold shock (Phadtare and Inouye 2004) were screened. A total of 55 (81%) showed altered morphology to cold shock (Knockouts presenting alterations to the cold shock response are reported together in Table 1, sorted by phenotype). The knockouts displaying altered response to cold shock represent a variety of cellular functions. Phadtare and Inouye report genes which respond to cold shock by altering their transcription levels. Comparisons (Chi^2^ not shown) between these transcription changes in *E. coli* and the cold shock phenotype displayed by these genes orthologs in Neurospora do not suggest there are any clear associations between transcription changes and cold shock morphology.

The screen of cold shock orthologs provides a test of the hypothesis that the cold shock response in both *E. coli* and Neurospora share a great deal of their cold shock response in common. The very high percentage of overlap between genes playing a role in these two widely separated organisms argues that the two responses are functionally related.

### Screen of Morphological/Hyphal plate knockout sets

A total of 129 selected mutant strains from the Neurospora knockout library were previously segregated into two collections. The “Morphological” collection resulted in known morphological variations in the knockout strains. The “Hyphal” collection consisted of knockouts of genes previously suspected to play a role in hyphal growth. These two collections were screened for alterations to their response to cold shock. In total, 30 (23%) strains were identified (Table 2) that displayed variant cold shock responses. The altered responses fell into several phenotypic categories (Table 1).

### Screen of transcription factor knockout set

A total of 147 Neurospora strains with knockouts in genes which function as transcription factors were screened for their response to cold shock. In all, 30 (20%) showed altered morphology to cold shock (Table 1).

As with the knockouts of orthologs of *E. coli* cold shock responding genes, the mutant strains identified in the additional screens show no observed correlations between the phenotypes observed and the annotated functions of the genes with a variety of functions being associated with the observed cold shock variations.

### Frequency of knockouts yielding alterations in the cold shock was dependent on the source of the knockout

As detailed above, mutants screened represented three different sets of knockouts: *E. coli* cold-shock responding orthologs, Neurospora morphological/hyphal growth mutants, and Neurospora transcription factors. These three groups displayed altered cold shock responses at different rates with the majority (81%) of the *E. coli* orthologs showing altered responses and much lower frequencies (23% and 20% respectfully) of the morph/hyphal and transcription factor knockouts showing altered responses (Table 1). Additionally, the phenotypes of the altered cold shock response showed a non-random distribution with regard to the knockout set the mutant was associated with using Chi^2^. Comparing knockout set vs cold shock phenotype among those with alterations yields a Chi^2^ of 32.2 and an associated p value < 1%. Much of the significance is coming from an over-representation of “dense” cold shock responses among otherwise unidentified (i.e. “hypothetical protein”) transcription factors.

### Cold shock phenotype was not correlated to GO categorization of the knockouts

The cold shock phenotype of knockouts was compared to their gene ontology categorizations via Chi^2^ analysis. Comparing cold shock phenotype to either its Molecular Function or Biological Process categorization failed to produce significant differences (p values of ~0.75 and ~0.5 respectfully). This fails to support the suggestion that knockouts with specific GO categories are associated with specific altered phenotypes in the cold shock response.

The data was also examined to determine if there was a non-random association between knockouts which show any alteration to their cold shock response (regardless of the specific phenotype) and those that show the wild type response vs their GO categorization. For both “Molecular Function” and “Biological Process” GO categories, no significant association was seen (via Chi^2^, p=0.4 and 0.5 respectfully), similarly failing to support the suggestion that knockouts with specific GO categorizations are tied to the cold shock response.

### Cold shock phenotype was weakly associated with growth rate among transcription factor knockouts

Linear growth rates for the transcription factor knockouts reported by Carrillo et al (2017) were compared via T-test for knockouts showing altered cold shock responses vs those showing no alteration to the response. One possible association between growth rate and altered cold shock phenotype was found for the knockouts displaying a dense phenotype which showed statistically faster growth rates than those with no alterations to cold shock (T-test, p=0.019). This is consistent with previous observations between growth rate and cold shock (Watters et al 2000), however the opposite association (slow growth rates among mutants displaying weak cold shock responses or failure to respond) is not observed, as would be expected if growth rate was a key factor among the knockouts. Taken together, there appears to be, at best, a weak association between growth rate and alterations to the cold shock phenotype among the transcription factor knockout mutants. This stands in contrast to the observation in wild type Neurospora (Watters et al 2000) that the morphology of the cold shock response was directly dependent on growth rate changes. This suggests that the altered morphologies observed among the knockout mutants are due to changes in gene activity associated with the knockouts and not simply the consequence of changes in growth rates in these mutants.

In conclusion, the gene functions highlighted by these screens (Table 1) are diverse. It is unclear how the diverse gene network, partially exposed here, coordinates for the function of temperature acclimatization. The results presented here demonstrate a strong relationship between the cold shock responses of *E. coli* and *Neurospora crassa*. The phenotype under examination here (morphological response to cold shock) appears to be influenced by a diverse network of genes. Similar diversity of function has been observed in other examinations of morphogenesis in Neurospora (Seiler & Plamann 2003). Further work on cold acclimatization should help clarify these connections.

